# Large language models help facilitate the automated synthesis of information on potential pest controllers

**DOI:** 10.1101/2024.01.12.575330

**Authors:** Daan Scheepens, Joseph Millard, Maxwell Farrell, Tim Newbold

**Affiliations:** University College London, UK; Natural History Museum London, UK; University of Glasgow, UK

**Keywords:** GPT-4, Information Synthesis, Large Language Models, Text Mining

## Abstract

The body of ecological literature, which informs much of our knowledge of the global loss of biodiversity, has been experiencing rapid growth in recent decades. The increasing difficulty to synthesise this literature manually has simultaneously resulted in a growing demand for automated text mining methods. Within the domain of deep learning, large language models (LLMs) have been the subject of considerable attention in recent years by virtue of great leaps in progress and a wide range of potential applications, however, quantitative investigation into their potential in ecology has so far been lacking. In this work, we analyse the ability of GPT-4 to extract information about invertebrate pests and pest controllers from abstracts of a body of literature on biological pest control, using a bespoke, zero-shot prompt. Our results show that the performance of GPT-4 is highly competitive with other state-of-the-art tools used for taxonomic named entity recognition and geographic location extraction tasks. On a held-out test set, we show that species and geographic locations are extracted with F1-scores of 99.8% and 95.3%, respectively, and highlight that the model is able to distinguish very effectively between the primary roles of interest (predators, parasitoids and pests). Moreover, we demonstrate the ability of the model to effectively extract and predict taxonomic information across various taxonomic ranks, and to automatically correct spelling mistakes. However, we do report a small number of cases of fabricated information (hallucinations). As a result of the current lack of specialised, pre-trained ecological language models, general-purpose LLMs may provide a promising way forward in ecology. Combined with tailored prompt engineering, such models can be employed for a wide range of text mining tasks in ecology, with the potential to greatly reduce time spent on manual screening and labelling of the literature.

## Introduction

Much of our knowledge of the global loss of biodiversity stems from large-scale syntheses of the ecological literature (Cornford et al., 2022). Such syntheses underlie the establishment of global environmental databases such as the WWF’s Living Planet Index (LPI, 2024), the PREDICTS (Hudson et al., 2017), and the BioTIME (Dornelas et al., 2018) databases, as well as the myriad global reports such as those of the Intergovernmental Science-Policy Platform on Biodiversity and Ecosystem Services (IPBES, 2019) and the Living Planet Report (Almond et al., 2022).

The body of ecological literature has simultaneously been experiencing rapid growth in recent decades (McCallen et al., 2019; Anderson et al., 2021) and it is becoming increasingly difficult to synthesise this literature manually (Cohen et al., 2012; Ananiadou et al., 2009). Thus, there is a growing demand in the ecological community for automated methods to assist in such tasks. Text mining and natural language processing (NLP) methods are expected to have significant potential in automating tasks such as document classification, named entity recognition and disambiguation, and the extraction of relations between entities (Farrell et al., 2022). Previous approaches have focused on the identification of species in the text using probabilistic machine learning algorithms (Akella et al., 2012) or dictionary-based approaches (Gerner et al., 2010); the extraction of taxonomic terms and geographic locations (Millard et al., 2020; Cornford et al., 2022); the extraction of taxonomic terms using deep learning (Le Guillarme and Thuiller, 2022); the extraction of population trends using random forest and neural network classifiers (Cornford et al., 2022); and the classification of relevant and non-relevant scientific articles using logistic regression and convolutional neural network approaches (Cornford et al., 2021).

Within the domain of deep learning (DL) for NLP tasks, large language models (LLMs) have been the subject of considerable attention in recent years by virtue of great leaps in progress and a wide range of potential applications (OpenAI, 2023a). This deep learning revolution in NLP is primarily fuelled by the now ubiquitous transformer architecture (Vaswani et al., 2023), which underlies many new and innovative DL tools in the natural sciences, such as the AlphaFold model for protein structure prediction (Jumper et al., 2021). Recent, transformer-based LLMs are trained on large amounts of input data to be able to generate realistic texts and are able to provide question answering and human-computer interaction via natural language (Ouyang et al., 2022). Provided with the right prompts, these models, furthermore, have been shown to exhibit advanced reasoning and problem-solving capabilities (Wei et al., 2023; Wang et al., 2023; Li et al., 2023; Kojima et al., 2023; Zhou et al., 2023). In particular, OpenAI’s fourth-generation Generative Pre-trained Transformer (GPT-4) has seen major improvements over previous models across a variety of benchmarks (OpenAI, 2023a). GPT-based models have already been investigated for a wide range of text mining tasks for research, including clinical (Hu et al., 2023), medical (Chen et al., 2023; Fink et al., 2023) and agricultural (Zhao et al., 2023), however, quantitative investigation into their potential in ecology has so far been lacking.

GPT-4 represents the current state-of-the-art of large language models and is easily accessed out of the box using ChatGPT, making this an attractive choice to demonstrate the potential of LLMs for automated information extraction and knowledge synthesis from scientific text. However, it is important to note that, as an alternative to closed access models such as OpenAI’s GPT series, the landscape of open-source LLMs is rapidly evolving and highly competitive (E.g., Touvron et al., 2023; Zhang et al., 2022; Dey et al., 2023).This availability of open-source LLMs is crucial for fostering open and reproducible use of AI in ecology, ensuring that the field advances in a sustainable and equitable manner. Thus, while GPT-4 does not currently adhere to open science standards, the methodology followed in this paper is to be understood as a proof of concept for the potential use of general-purpose LLMs in information extraction tasks in ecology, which ideally transpire in an open science context.

This work investigates the application of GPT-4 to a body of ecological literature on biological pest control. The utilisation of natural enemies of pests, such as arthropod predators and parasitoids, as biological control agents can provide an effective way to reduce pesticide usage, which is currently a major driver of insect declines (Wagner et al., 2021; Sánchez-Bayo and Wyckhuys, 2019; Cardoso et al., 2020). Biological control has historically often relied on the introduction of non-native species (classical biocontrol), which can be detrimental to native species and thus strain local ecosystems and negatively affect biological diversity. Natural biological control, on the other hand, utilises native species as biological control agents and is typically achieved through the enhancement of natural habitat. Natural biological control helps directly regulate the frequency of pest outbreaks (Letourneau, 2012; Tahvanainen and Root, 1972; Pimentel, 1961) and, indirectly, can result in improved soil quality (Gunstone et al., 2021), increased crop yields (Gurr et al., 2016; Dainese et al., 2019; Letourneau et al., 2011) and increased abundances of other beneficial organisms such as pollinators (Balzan et al., 2014; Grass et al., 2016; Wratten et al., 2012), as a result of reduced pesticide usage and increased natural habitat. Crucially, by strengthening the stability and resilience of ecosystem services such as pest control, pollination and nutrient cycling, food production systems can be better buffered against environmental and climatic changes (Oliver et al., 2015; Brittain et al., 2013; Martin et al., 2019), which is of growing importance as the impacts of climate change continue to intensify (IPCC, 2023).

The primary aim of this paper is to provide a thorough analysis of the ability of GPT-4 to reliably extract information from scientific abstracts to identify pests and pest controllers. In addition to determining roles of species (e.g., as pests or pest controllers), the model is tasked with extracting their taxonomy and geographic location, as well as role-specific information such as the pest-type and the crop or plant that a pest affects. As such, the task comprises multiple sub-tasks, which require both the capability to recognise and disambiguate entities (named entity recognition), and to extract relations between entities (relation extraction). Rather than fine-tuning the parameters of the model, we proceed to optimise model performance through the fine-tuning of the prompt itself. Performance is then analysed on each sub-task of the query, indicating precision, recall and F1 scores. To our knowledge, this is the first instance of general-purpose, large language models such as GPT-4 being investigated for the potential automation of information extraction and knowledge synthesis in ecology.

## Methods

### Data collection

In order to obtain relevant literature on potential pest controllers and their hosts, we extracted a set of abstracts from the academic indexing tool Scopus up until the year 2020, using the following search-term:

> TITLE-ABS-KEY(“pest control” OR “biological control” OR “pest management” OR “natural enem*”) AND (LIMIT-TO(DOCTYPE, “ar”)) AND (LIMIT-TO(SUBJAREA, “AGRI”) OR LIMIT-TO(SUBJAREA, “ENVI”)) AND (LIMITTO(LANGUAGE, “English”))

The usage of Scopus ensured that all of the extracted literature has undergone peer-review, which we judged to be of importance, given that the underlying motive of this work is to automate peer-reviewed knowledge synthesis and meta analysis. The search-term used here obtained a corpus of 58,791 abstracts. For this study, we selected a subset of 100 abstracts to create a training set, which we used to fine-tune the prompt with respect to GPT-4 output, and a further 100 abstracts to create a held-out test set, which we used to analyse the final performance of GPT-4 with the fine-tuned prompt. We populated both of these sets by manually labelling the 200 abstracts (including titles and keywords) using the columns shown in Table 1, including all invertebrate species found at the species or genus level. We made this selection of species due to our focus on pest control services of, and provided by, invertebrate animals specifically. Invertebrates also comprise the vast majority of species present in the abstracts selected for the training set and the held-out test set.

**Table 1.** Columns of the data used in this study. Using these columns, we manually populated the training set (using the 100 abstracts selected for the training set), and the test set (using the 100 abstracts selected for the test set): These constitute the ‘manually labelled data’ in the training and test set. Consequently, we instructed GPT-4 to generate a table with the same columns: These constitute the ‘GPT-4 generated’ training and test set.

| Column | Explanation | Clarification |
| --- | --- | --- |
| 1. Class | Class name. | Taxonomy (Latin). |
| 2. Order | Order name. | Taxonomy (Latin). |
| 3. Family | Family name. | Taxonomy (Latin). |
| 4. Genus | Genus name. | Taxonomy (Latin). |
| 5. Species | Species name. | Taxonomy (Latin). |
| 6. Role | The role that the species is ascribed in the text (e.g., predator, parasitoid, pest, herbivore, etc.). | Not limited to the given examples. |
| 7. G/S | Is the species mentioned as a generalist or specialist? | This refers to dietary generalism and specialism. |
| 8. Pest Controller | True or false: Is the species mentioned as a pest controller? | We define a pest controller as a natural enemy (i.e., predator or parasitoid) of a pest. Biocontrol agents of pests also suffice. |
| 9. Pest Names | For pest controllers: What pests does it control? | May be given as a common name or at any taxonomic scale, as mentioned in the abstract. |
| 10. Pest Type | For pests and pest controllers: Is the pest an invertebrate or plant (i.e., weed)? | Although plant species are not to be returned as species in the table, they may be mentioned in Column 9. |
| 11. Industry Type | For pests and pest controllers: What industry is the pest associated with (e.g., agriculture, forestry, freshwater, etc.)? | Not limited to the given examples. |
| 12. Affects | For pests and pest controllers: What crops, plants or products does the pest affect? | This may be given as common names or at any taxonomic scale, as mentioned in the abstract. |
| 13. Description | Thorough description of the species as mentioned in the abstract and a justification for the assessment. | Unconstrained in length. |
| 14. Location | Geographical location of the study, if mentioned. | Name of the location as mentioned in the abstract, unconstrained in scale. |

We selected the 100 abstracts comprising the training set on the basis of specifically including both predators and parasitoids, in addition to pests, in order to fine-tune the prompt on these roles as much as possible, as these roles bear particular relevance to the topic of (natural) biological pest control. The information on instances of predators, parasitoids and pests was available from a pre-screening procedure that was carried out in previous work for a total of 1520 abstracts, which identified and described genera present in these abstracts and could thus indicate which abstracts contained (at least one genera) of predators, parasitoids and pests. The abstracts comprising the training data were then selected from this subset at random on a rolling basis, with an ongoing attempt to avoid extreme imbalances between the number of predators, parasitoids and pests (Fig. S1.1a). The 100 abstracts comprising the held-out test set were selected randomly from the remaining abstracts in the corpus. Pest species are common in the corpus, which means that there is a relative increase in the proportion of species labelled as pests in the test set compared to the training set (Fig. S1.1b).

The training and test set consist of 14 columns (Table 1). In the ‘Role’ column, we determined whether a species was mentioned as a predator, parasitoid or pest. If there was no mention of these roles in the text, we looked for any other broad category, such as herbivore, pollinator, biological control agent, etc. Where specific roles lacked, we looked for broader descriptions, such as “was observed to predate/consume”, “was reared from host” or “infests maize fields” to arrive at an appropriate role. Where even descriptions lacked and no other indication was given in the abstract, the role was left blank. In the ‘Generalist/Specialist’ column, we determined whether a species was mentioned as a (dietary) generalist or specialist. We required no specific evidence for this column, only the mention of dietary generalism or specialism. For the ‘Pest Controller’ column to be ‘true’, we required either a specific mention of a species as functioning as a biological control agent, or required a predator, parasitoid or natural enemy to be stated to specifically feed or prey on a pest. In the ‘Pest Names’ column, we specified the names of the pests controlled by a pest controller. In the ‘Pest Type’ column, we decided to distinguish invertebrate pests from plant pests (weeds), with the aim of identifying weed controllers from other pest controllers. In the column ‘Industry Type’ we determined if a pest species was mentioned in association with a particular industry like agriculture or forestry, and in the column ‘Affects’ we determined which crops or products were affected by the pest, if mentioned in the abstract. The column ‘Description’ serves to provide both as a description of the species in the abstract, and as a justification for the assessment of the other columns. The ‘Location’ column refers to the location of the study, if mentioned, and was unconstrained in geographical scale.

### Prompt and objective

In the first step of the experiment, we applied GPT-4 to each individual abstract (title, abstract and author keywords) in the training set, with the instruction to find all taxonomic entities present in the text that are mentioned at the speciesor genus level and returning a table with columns as in Table 1. To achieve this, we instructed GPT-4 with an initial prompt (Fig. S3.1). This prompt follows a zero-shot prompting approach, detailing precisely how each column in the table should be filled out, requiring both named entity recognition and relation extraction capabilities.

Since the training and validation set were manually populated with invertebrate species only, we decided to prompt GPT-4 to neglect plants, bacteria, fungi and pathogens in its output. While this does not exclude vertebrate species per se, we proceed to analyse the performance of the model on invertebrates only. The ability of GPT-4 to abide by the constraint to neglect certain entities in the text will be of interest in cases where generated output is desired to be limited to certain species of interest, rather than providing an exhaustive table. For large-scale studies, the increased completion time of generating exhaustive tables may be substantial, and the generation of long tabular output may be infeasible due to the token limits posed by current state-of-the-art large language models such as GPT-4. At the time of this experiment, both the GPT-4’s input and generation were limited to 2,048 tokens (OpenAI, 2023b), corresponding to ca. 1536 words (OpenAI, 2023c).

As GPT-4 had not yet been released for API usage at the time of the study, we proceeded to use the ChatGPT web-interface (using the May 24 release of GPT-4) and saved the tabular output manually to a spreadsheet. This required a Plus membership with OpenAI. We prompted the model with each abstract in an individual session to ensure that there was no information leakage across abstracts. We then analysed the output for each abstract in the training set, as obtained with the initial prompt (Fig. S3.1), noting its errors across the various columns of the table, and subsequently improved the prompt through a series of changes that proved to remedy the error in each particular case. This was achieved through a combination of experimenting with the prompt design, based on consultation of the prompt learning literature, and inserting bespoke instructions and clarifications into the prompt (e.g., “Don’t do *X* “, “Ensure that *Y* “). Where uncertain how to proceed, we queried GPT-4 itself as to how the prompt may be improved in order to avoid a particular error in the future. This was done by following up on the output with “You made the following error: *X*. How would I need to change my prompt in order for you to fill this out correctly next time?”. We carried out this fine-tuning process for the 100 abstracts in the training data over the course of three full iterations. We decided to iterate over the training set three times, rather than once, in order to ensure that the continuous changes made to the prompt yielded consistent results and to allow for changes to be reverted if they proved ineffective. All consecutive versions of the prompt over the course of the fine-tuning process can be found in Section 3 of the supplementary information.

Following fine-tuning against the training set, the prompt assumed the design as shown in Fig. 1. This prompt is split into three smaller prompts, the first of which serves to identify all relevant species in the abstract, the second of which serves to return these species in a table with the appropriate columns filled out, and the third of which asks GPT-4 to review the generated table and make any corrections if necessary. We ran these three prompts within the same session. This approach, of dividing a complex set of instructions into smaller tasks, is known as least-to-most prompting (Zhou et al., 2023). The prompt design follows a ‘zero-shot’ approach as it contains only instructions, rather than including any exemplar prompt completions, which would have drastically increased the token length of the prompt. We attempted to improve the reasoning ability of the model by including the phrase “let’s think through the following tasks step by step” (info-point 3; Kojima et al. (2023)), and through the usage of chain-of-thought reasoning (Wei et al., 2023) at several points in the prompt to demonstrate clearly to the model how it should think about its tasks and what it must pay attention to (infopoints ‘C’). These prompting techniques have all been demonstrated to improve the reasoning capabilities of GPT-3.5 (Zhou et al., 2023; Kojima et al., 2023; Wei et al., 2023). The prompt also makes abundant usage of specific examples and counter-examples to aid the correct identification of particular roles of entities. We applied this fine-tuned prompt to the training data once more without any further changes in order to obtain final results on the training set, and subsequently applied the prompt to the held-out test set.

**Figure 1.**
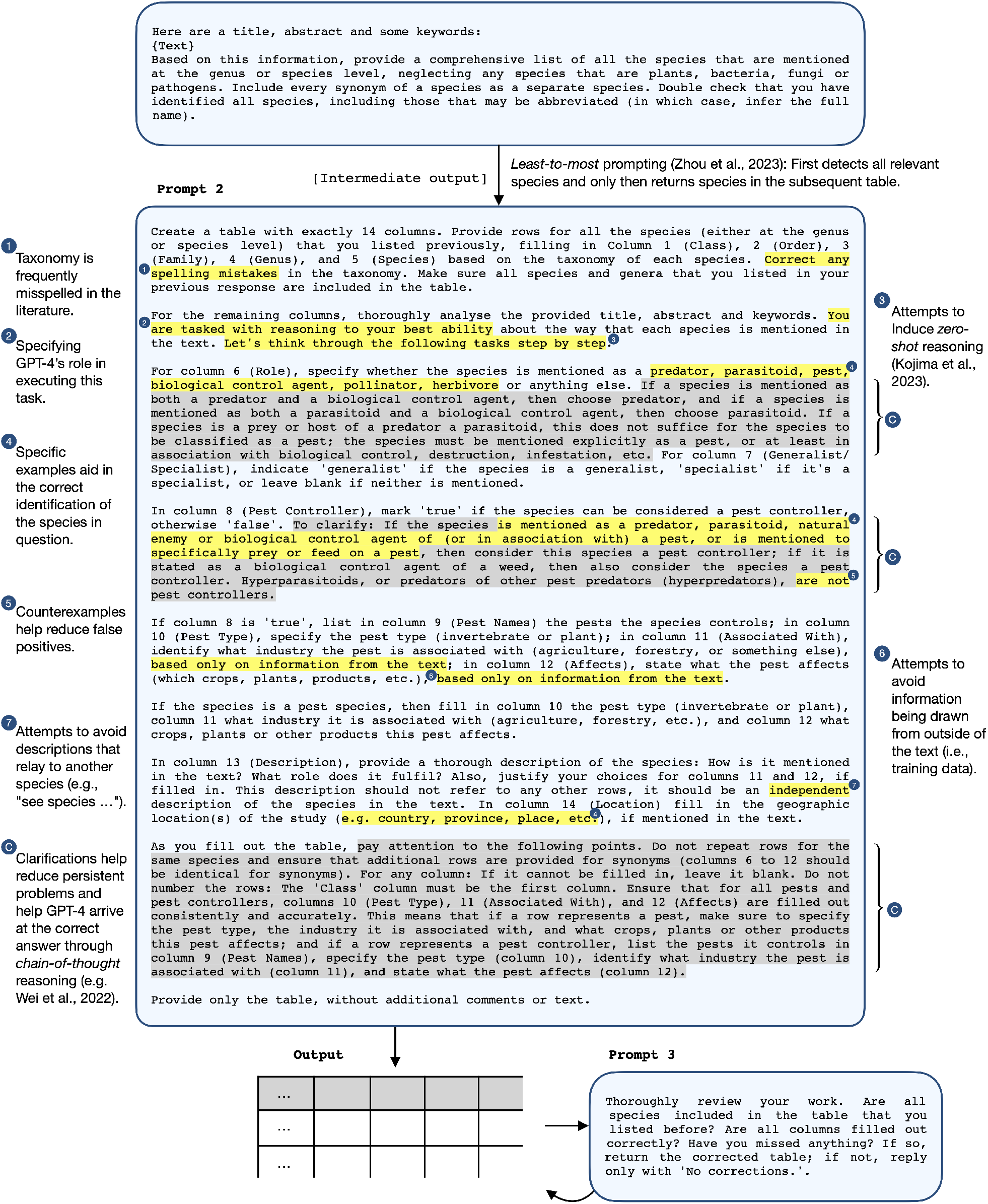
The final prompt design after fine-tuning against the training set. The prompt is divided into three parts. The first part prompts GPT-4 to detect all relevant species in the abstract, the second part prompts GPT-4 to return these in the table with the appropriate columns filled out, and the third part prompts GPT-4 to review the generated table and make any corrections if necessary. The points labelled as ‘C’ (and highlighted in grey rather than yellow) designate clarifications; these were used to clarify specific terms such as pest and pest controller, and aimed to reduce persistent problems such as formatting errors and missing pest information in some of the columns.

### Extraction of taxonomy

As shown in Table 1, the first 5 columns of the data reflect the taxonomy of the species or genus (i.e., Class, Order, Family, Genus, Species). While higher level taxonomic information can be extracted from databases such as the Global Biodiversity Information Facility (GBIF, 2023) for a given species or genus, species names are frequently misspelled in the literature and genus names occasionally appear across different phyla, making it difficult to automatically extract the correct taxonomy. We thus deemed it of interest to investigate the ability of GPT-4 to (1) predict the missing taxonomy of a given species or genus (i.e., from its training data) and (2) correct any obvious misspellings. To this end, we filled out the taxonomy of each species (or genus) in the training and test sets as they are stated in the abstracts, if available. We refer to the task of extracting this taxonomy as ‘taxonomic named entity recognition’. Where the taxonomy was not available in the abstract, we searched for the species on GBIF and the Encyclopedia of Life (EOL) (Parr et al., 2014) using the genus and (if available) species name and filled this out accordingly. We refer to the task of predicting this missing taxonomy as ‘higher level taxonomy prediction’. This taxonomy concerns only the class, order and family columns in the data sets as the genus and species columns are needed to predict the missing taxonomy. In both cases, the obtained taxonomic information is then compared to the generated GPT-4 output. Since taxonomic named entity recognition and higher level taxonomy prediction comprise two very different tasks, we evaluate performance of the model on these two groups of taxonomic terms individually.

#### Mismatches

A mismatch between the manually labelled taxonomy and the taxonomy returned by GPT-4 does not necessarily mean that the GPT-4 prediction is wrong. Taxonomy is frequently misspelled in the literature and conventions regarding taxonomic ranks are not always used consistently. For example, Latin binomials are often accompanied by a taxonomic annotation ‘(X: Y)’, where X typically refers to the order and Y typically refers to the family of the species. However, authors frequently present this annotation with phyla, class, suborders or superfamilies instead, which means that the manual extraction does not always extract the correct class, order and family information. In order to determine the correct taxonomic ranks in case of a mismatch between the term manually extracted from the abstract and the GPT-4 generated term, we once again referred to the GBIF and EOL databases. Through comparison with this reference taxonomy, we then determined whether the manual label, the GPT-4 prediction, or both, were mistaken. Conversely, a mismatch between taxonomy that was missing from the abstract and the taxonomy predicted by GPT-4 necessarily implies that the GPT-4 prediction was mistaken, since the missing taxonomy was obtained from the GBIF and EOL databases to begin with.

We proceed to divide the taxonomic mistakes made by GPT-4 into two categories: Minor mistakes and major mistakes. We define a minor mistake as a discrepancy between the GPT-4 prediction and the manual label where the GPT-4 prediction nevertheless preserves essential details. This includes predictions of incorrect taxonomic ranks that are nevertheless part of the broader taxonomy as adjacent terms (e.g., the correct suborder rather than order, phylum rather than class, etc.), entries of ‘sp’/’spp.’ rather than blank species entries, inclusions of subspecies information, or common names rather than Latin names (e.g., ‘Insect’ rather than ‘Insecta’). Conversely, we define major mistakes as discrepancies where the GPT-4 prediction conveys information that is inaccurate. This refers to incorrect taxonomic terms that are not part of the broader taxonomy of the species, and instances of blank entries where the taxonomy is stated in the abstract. We refer to approximate taxonomic matches as matches that include both exact matches and minor mistakes.

### Analysis of species information

We focus our analysis of the remaining species information (columns 6–14) on the species roles and the geographic locations. For these two sub-tasks, we measure the performance of GPT-4 by computing the precision, recall and F1-score (Eq. 1) of the model, and provide confusion matrices to identify true positives, false positives, true negatives and false negatives for each label. The F1-score serves to strike a balance between the precision and the recall.

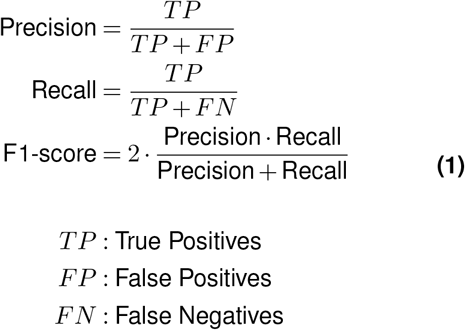

True positives comprise all direct matches between the manual labels and the GPT-4 predictions, as well as predictions in close agreement with the manual label (i.e., convey the same essential information). For species roles, this is achieved by grouping all obtained roles into 9 distinct groups, with which we ensure that roles such as ‘biocontrol agent’ and ‘biological control agent’ both comprise the same essential role. Similarly, roles such as ‘parasitoid’ and ‘larval parasitoid’ comprise the same group, as do labels such as ‘unclear’, ‘not mentioned’ and blank entries. We group roles that are not of immediate interest to this study as ‘other’ (e.g., ‘pollinator’, ‘scavenger’, ‘competitor’).

For geographical locations, ‘close agreement’ between labels can be understood as referencing the same geographic area, albeit at different scales. As such, we count examples such as ‘Northern New Zealand’ and ‘New Zealand’, ‘Manitoba, western Canada’ and ‘Manitoba, Canada’, ‘Eastern Newfoundland’ and ‘Newfoundland’ (taken from the training set) as the same essential location. Naturally, when the essential location is lost (e.g., a manual label of ‘Australia’ and a prediction of ‘North Island’) we no longer consider this a true positive (indeed, this example would comprise a false negative, as it missed the essential location, Australia).

## Results

### Species extraction

We found that splitting the instructions into two consecutive prompts, in which the first prompt served to identify all relevant species in the text and the second prompt served to fill out the table, led to improvement in entity extraction capability. While the initial prompt already extracted 627 out of the 649 species (96.6% recall) in the training data (with missing entries corresponding to 2 out of 100 abstracts), the split-prompt approach ensured that all 649 species were extracted in the first part of the prompt. In the second part of the prompt, however, GPT-4 remained challenged with returning all previously found species in the final table, and as a result a small number of species in the training data were still missed in GPT-4’s output, with 631 out of 649 species (97.2% recall) extracted (Table 2a). However, since these missing entries correspond to only 2 out of 100 abstracts in the training set, we obtain a mean recall per abstract of 99.5% (standard deviation = 4.4%) (Table 2b). Importantly, we also observed 43 instances of fabricated species entries (‘hallucinations’) as rows in the GPT-4 output for the training set, which occurred across the same two abstracts where missing entries were observed. This results in a total precision of 93.6% and a mean precision per abstract of 99.1% (standard deviation = 6.8%) (Table 2). For the test set, GPT-4 extracted 244 out of 245 species (99.6% recall), and we observed no hallucinated species entries in the test set (100% precision) (Table 2). Moreover, we found that GPT-4 did not generate any entries for plants, bacteria, fungi or pathogens in either the training set or the test set and thus managed to abide by the constraint set on the species extraction very effectively.

**Table 2.** Precision, recall and F1-score of the extraction of species (as rows in the final output table) by GPT-4, for the training set and the held-out test set. Scores are presented as computed from the total number of predicted and manually extracted species entries (a), and as the mean per abstract (b). Support: The number of species present in the manually labelled data set (a); the number of abstracts in the data set (b).

| (a) Total |  |  |  |  |
| --- | --- | --- | --- | --- |
|  | Precision (%) | Recall (%) | F1 (%) | Support |
| Training set | 93.6 | 97.2 | 95.3 | 649 |
| Test set | 100.0 | 99.6 | 99.8 | 245 |

| (b) Mean per abstract |  |  |  |  |
| --- | --- | --- | --- | --- |
|  | Precision (%) | Recall (%) | F1 (%) | Support |
| Training set | 99.1 ± 6.8 | 99.5 ± 4.4 | 99.3 ± 4.1 | 100 |
| Test set | 100.0 ± 0.0 | 99.0 ± 9.9 | 99.5 ± 5.0 | 100 |

We hypothesised that the third step in the final prompt design, in which we asked GPT-4 to thoroughly review its output, would allow GPT-4 to pick up on its own mistakes, however, its success was found to be very limited. GPT-4 corrected its own mistakes only in the case of a single abstract in the training set, corresponding to 12 species entries. In this case, the model correctly identified that it had left out columns 10 (‘Pest Type’), 11 (‘Industry Type’) and 12 (‘Affects’) for the pest species in the table and then proceeded to fill these out correctly. In one case in the test set, the model claimed that it had not identified a particular species as ‘pest’, stating that it had corrected this, although the species had, in fact, been identified as a pest already. In two other cases, the model returned reassurances other than ‘No corrections’ (although conveying the same message). Interestingly, the model did not recognise missing entries or hallucinations in the table.

The generated tables obtained from GPT-4 display varying degrees of accuracy and adherence to the prompted instructions. However, in cases where the information in the abstract is presented clearly and non-ambiguously, strong performance can be observed (Fig. 2). Conversely, ambiguous usage of language (e.g., a predatory species that acts as a pest) is prone to produce erroneous results (Fig. S1.2). The results in the following sections are obtained from comparing the GPT-4 generated tables (using the fine-tuned prompt; Fig. 1) with the manually labelled tables for all abstracts in the training and the held-out test set. The species in the training and test set that were missed by GPT-4 are omitted from the remaining analysis as these species naturally cannot be compared to GPT-4 predictions. This refers to 18 out of 642 entries in the training set and 1 out of 245 entries in the test set. We also omit the 43 cases of hallucinations in the training set from this further analysis, but discuss these in the discussion.

**Figure 2.**
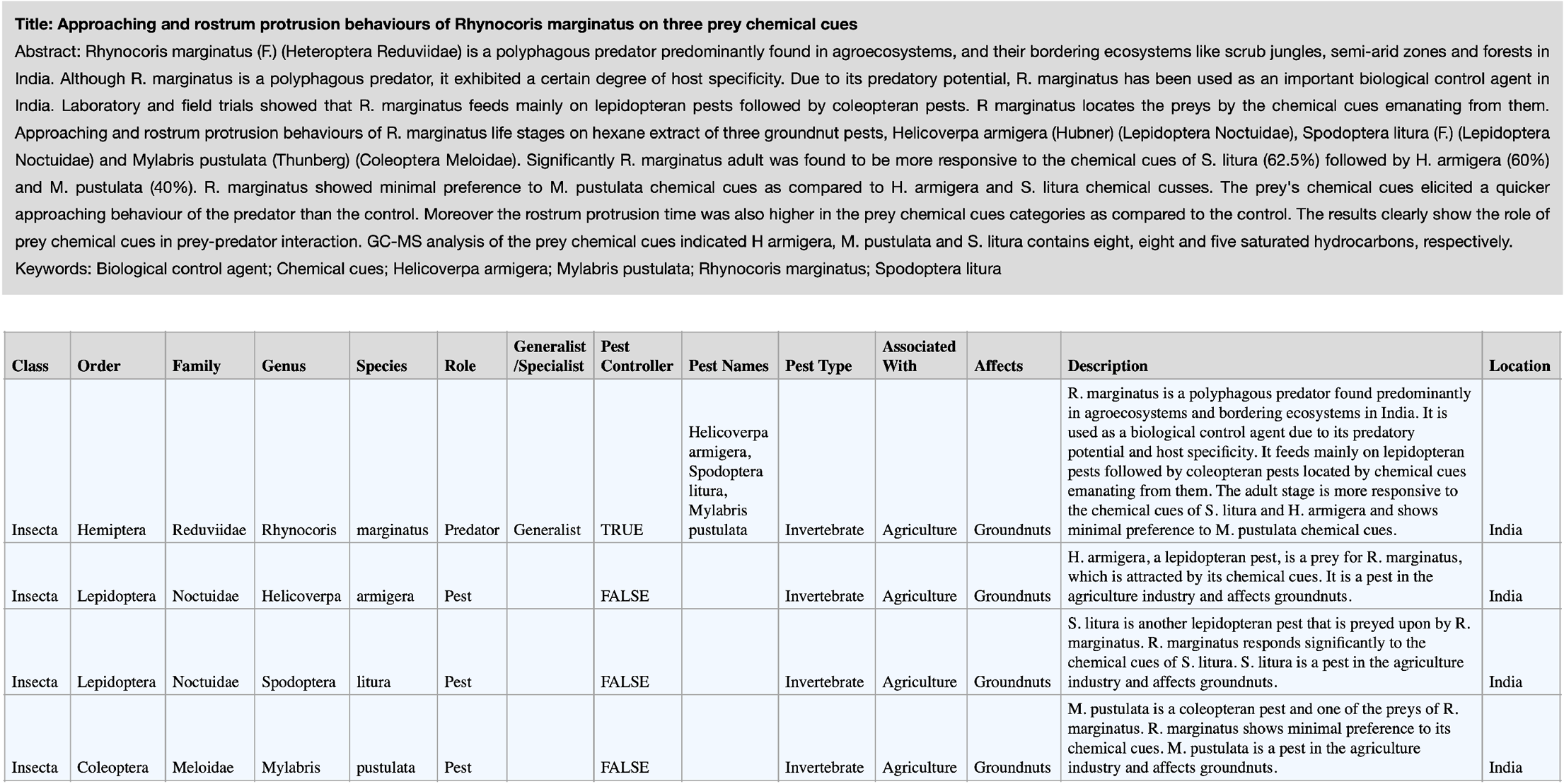
Table generated by GPT-4 for an exemplar abstract (title, abstract and keywords) in the test set, using the fine-tuned prompt design (Fig. 1). This is an abstract discussing a predatory true bug found in India (Sahayaraj, 2008). The generated table includes all four species mentioned in the abstract and correctly identifies R. marginatus as pest controlling predator and the remaining species as pests. The taxonomy is correctly extracted from the text, with the class correctly predicted to be ‘Insecta’. The order of R. marginatus has been correctly predicted to be Hemiptera rather than the suborder Heteroptera that is stated in the text. The description of R. marginatus as “polyphagous predator” is translated into ‘Generalist’ in column 7, although the “certain degree of host specificity” is perhaps neglected. The pest information (columns 9–12) is correctly filled out both for R. marginatus as pest controller and for the remaining species as pests. The descriptions are thorough and factually correct and the location is correctly identified to be India for all four species.

### Taxonomic named entity recognition

Here we assess the ability of GPT-4 to extract taxonomic information present in the abstracts. We observed only a small number of mismatches between the manual labels and the GPT-4 predictions (Table 3). Following comparison with the reference taxonomy obtained from GBIF and EOL, we found that the majority of mismatches were either cases where the GPT-4 prediction was, in fact, correct, or where the mistake comprised only a minor mistake, rather than a major mistake (Table S2.3a). Indeed, we observed only 2 major mistakes out of the 2276 total taxonomic terms (0.09%) predicted by GPT-4 in the training set, and 3 major mistakes out of the 803 total taxonomic terms (0.37%) predicted by GPT-4 in the test set (Table S2.3a). Proportion correct (PC) scores of approximate matches are found to be between 98–100% across both the training and test set, whether computed across all taxonomic terms S2.1). (PC_approx_) or across abstracts 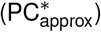 (Table 3).

**Table 3.**
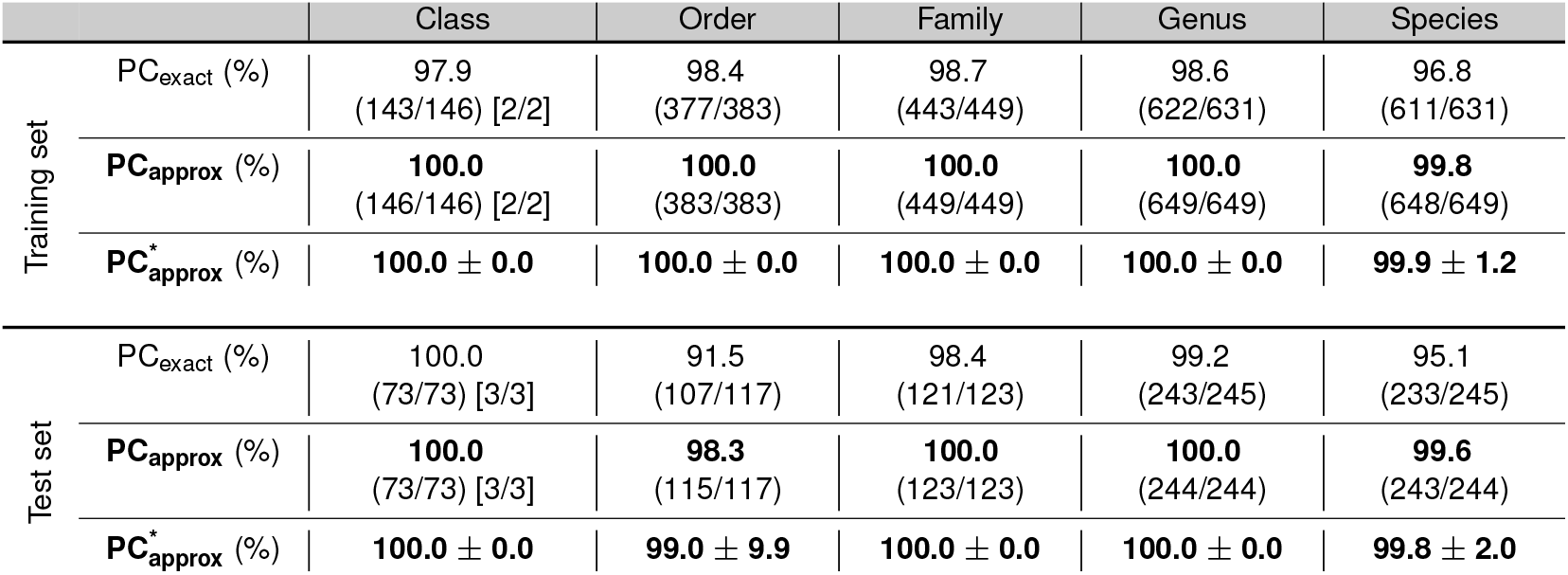
Accuracy of GPT-4 on the task of taxonomic named entity recognition. This refers to taxonomy that was available for extraction in the abstract. Accuracy is presented for each taxonomic rank as the proportion correct (PC) of predictions that are exact matches versus the total number of taxonomic terms (PC_exact_), and as the proportion correct of predictions that are approximate matches (which exclude major mistakes but retain minor mistakes) versus the total number of taxonomic terms (PC_approx_). We also computed the latter *per-abstract* 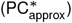 to avoid bias from abstracts with a large number of extracted species: The presented values show the mean across abstracts with a spread of one standard deviation. Since the class column in both the training and the test set is heavily dominated by insect species, we also provide information on the correct number of non-insects obtained in square brackets.

Mismatches in the genus and species columns where the GPT-4 prediction was correct refer primarily to cases of spelling corrections, while mismatches in the class, order and family columns refer primarily to cases of corrected ranks (e.g., the correct family as opposed to the superfamily that was mentioned in the abstract), although we also find spelling corrections in the family column. Spelling corrections, which occurred exclusively in the family, genus and species ranks, comprised 12 cases in the training set and 4 cases in the test set. In all cases, GPT-4 corrected the spelling mistake and returned the correct term. While this does not evidence that the all misspellings in the data were corrected by the model, it does demonstrate that when a misspelling was addressed, it was done correctly.

### Higher level taxonomy prediction

For the task of higher level taxonomy prediction we observed more mismatches than for taxonomic named entity recognition (Table 4), and more of these comprised major mistakes (Table S2.3b). We observe 15 major mistakes out of the 915 total taxonomic terms (1.64%) predicted by GPT-4 in the training set, and 7 major mistakes out of the 419 total taxonomic terms (1.67%) predicted by GPT-4 in the test set (Table S2.3a). While the PC scores of approximate matches (PC_approx_) can be seen to lie between 98–100% for the class and order, we observe a notable decline for the family rank in both the training set and the test set (Table 4). This decline, however, appears to be primarily a result of bias towards a small number of abstracts in which many major mistakes were made, and is thus not observed in the scores per abstract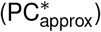.

**Table 4.**
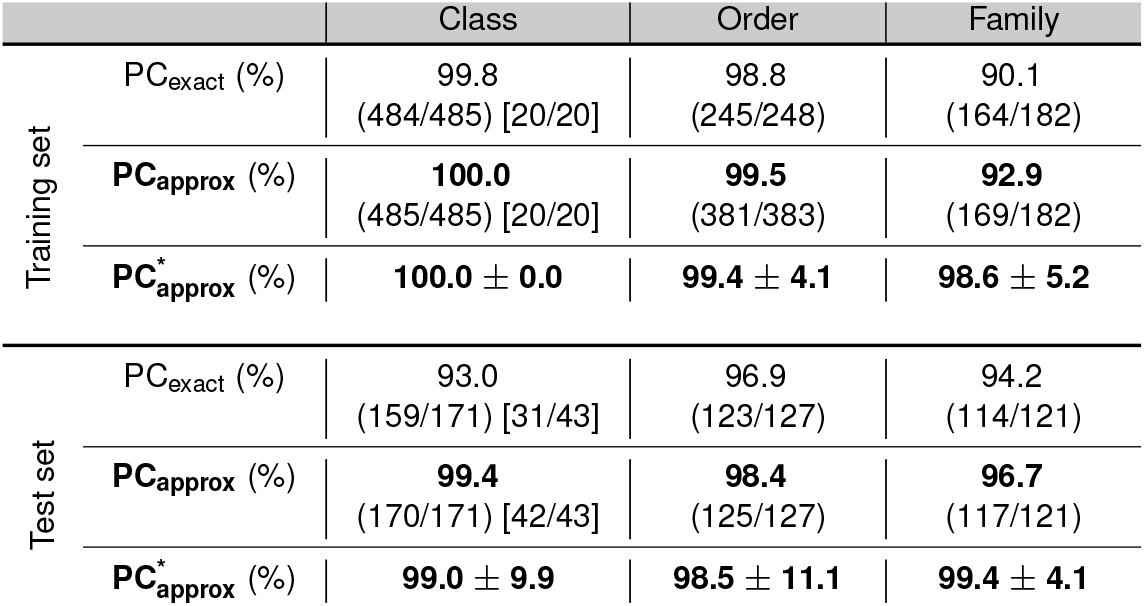
Accuracy of GPT-4 on the task of higher level taxonomy prediction. This refers to taxonomy that was not stated in the abstract and thus had to be predicted by GPT-4 based on its training data. Accuracy is presented for each taxonomic rank as the proportion correct (PC) of predictions that are exact matches versus the total number of taxonomic terms (PC_exact_), and as the proportion correct of predictions that are approximate matches (which exclude major mistakes but retain minor mistakes) versus the total number of taxonomic terms (PC_approx_). We also computed the latter *per-abstract* 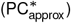 to avoid bias from abstracts with a large number of extracted species: The presented values show the mean across abstracts with a spread of one standard deviation. Since the class column in both the training and the test set is heavily dominated by insect species, we also provide information on the correct number of non-insects obtained in square brackets.

### Species Roles

GPT-4 captured the roles of species with a high degree of accuracy, as assessed against both the training set (Fig. 3a) and the test set (Fig. 3b). Crucially, while there are a small number of cases of manually labelled predators, parasitoids and pests having been mislabelled by GPT-4 as something else (e.g., as ‘competitor’, ‘prey’, ‘host’, ‘unclear’), there is no confusion between these roles, either in the training set, or in the test set (emphasised by the dotted line).

**Figure 3.**
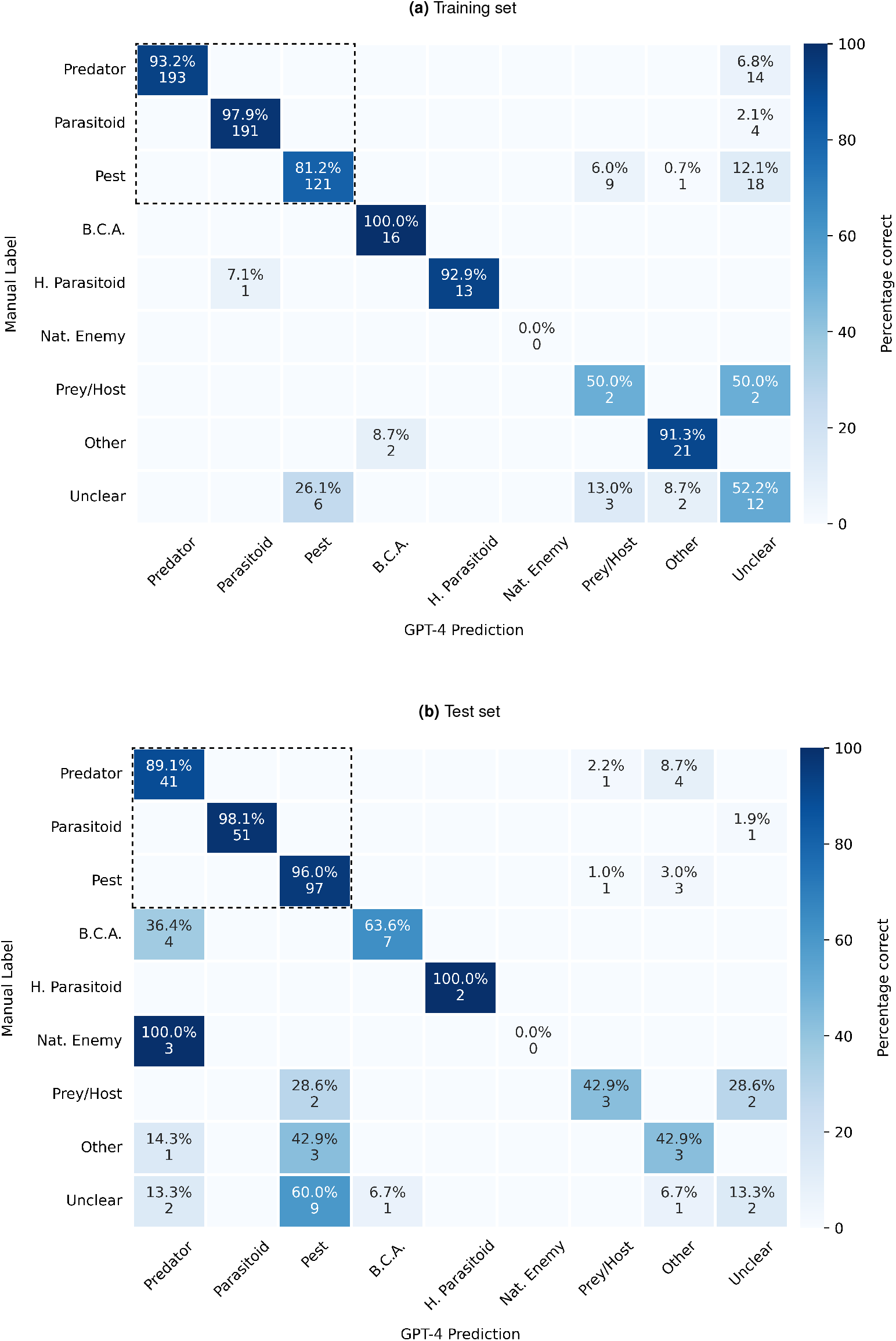
Confusion matrix of the various roles ascribed to species, either as ‘Manual label’ or as ‘GPT-4 prediction’, for the training set. Values indicate the percentage of total predictions across each row (recall) and below these the corresponding absolute numbers of predictions. The primary roles of interest (predator, parasitoid and pest) are emphasised with a dotted line. ‘B.C.A.’: Biological Control Agent. ‘H. Parasitoid’: Hyperparasitoid. ‘Nat. Enemy’: Natural enemy. ‘Other’ includes terms such as ‘Pollinator’, ‘Herbivore’, ‘Leaf miner’, ‘Scavenger’, ‘Ectoparasite’ and ‘Competitor’ and the term ‘Unclear’ includes ‘Not mentioned’, as well as empty entries.

Results on the test set are very comparable with results on the training set, although some differences in precision and recall can be observed (Table 5b). In particular, the precision for predator roles is reduced from 100% on the training set to 80.4% on the test set. As we can see from Fig. 3b, this reduction is primarily a result of GPT4 predictions of ‘predator’ for terms that were manually labelled as biological control agents, natural enemies or were left blank (unclear). Recall of predator roles is only marginally reduced on the test set as compared to the training set. In both cases, false negatives are a result of a comparatively small percentage of abstracts (Table Performance for parasitoid roles on the test set matches performance on the training set closely, with precision and recall scores consistently between 97–100% and an F1-score of 98.7–99.0% (Table 5). In both sets, hyperparasitoids (parasitoids of other parasitoids) are accurately distinguished. For pest roles, we observe a substantial amount of confusion between pests and ‘prey/host’, ‘other’ and (in particular) ‘unclear’, across both the training and the test set (Fig. 3a and Fig. 3b). We also observe a large variation (between the training set and the test set) in precision and recall for this role (Table 5).

**Table 5.** Precision and recall obtained by GPT-4 for each role in the training set (a) and the held-out test set (b). Support: The number of instances of the respective role in the manual labels. Weighted average: The average of the respective column, weighted by the support of each row. ‘B.C.A.’: Biological Control Agent. ‘H. Parasitoid’: Hyperparasitoid. ‘Nat. Enemy’: Natural enemy. ‘Other’ includes terms such as ‘Pollinator’, ‘Herbivore’, ‘Leaf miner’, ‘Scavenger’, ‘Ectoparasite’ and ‘Competitor’ and the term ‘Unclear’ includes ‘Not mentioned’, as well as empty entries.

|  | Precision (%) | Recall (%) | F1 (%) | Support |
| --- | --- | --- | --- | --- |
| **Predator** | **100.0** | **93.2** | **96.5** | 207 |
| **Parasitoid** | **99.5** | **97.9** | **98.7** | 195 |
| **Pest** | **95.3** | **81.2** | **87.7** | 149 |
| B.C.A. | 88.9 | 100.0 | 94.1 | 16 |
| H. Parasitoid | 100.0 | 92.9 | 96.3 | 14 |
| Nat. Enemy | NA | NA | NA | 0 |
| Prey/Host | 14.3 | 50.0 | 22.2 | 4 |
| Other | 87.5 | 91.3 | 89.4 | 23 |
| Unclear | 24.0 | 52.2 | 32.9 | 23 |
| Weighted average | 94.7 | 90.2 | 92.0 | 631 |

(b) Test set
|  | Precision (%) | Recall (%) | F1 (%) | Support |
| --- | --- | --- | --- | --- |
| **Predator** | **80.4** | **89.1** | **84.5** | 46 |
| **Parasitoid** | **100.0** | **98.1** | **99.0** | 52 |
| **Pest** | **87.4** | **96.0** | **91.5** | 101 |
| B.C.A. | 87.5 | 63.6 | 73.7 | 11 |
| H. Parasitoid | 100.0 | 100.0 | 100.0 | 2 |
| Nat. Enemy | NA | 0.000 | NA | 3 |
| Prey/Host | 60.0 | 42.9 | 50.0 | 7 |
| Other | 27.3 | 42.9 | 33.3 | 7 |
| Unclear | 40.0 | 13.3 | 20.0 | 15 |
| Weighted average | 82.4 | 84.4 | 82.7 | 244 |

### Geographic locations

In addition to the roles of species, we investigated the ability of GPT-4 to extract geographical information from the abstracts. The results show that GPT-4 is able to effectively extract geographic information, with few false-positive and false-negative predictions for both the training set and test set (Fig. 4). Indeed, locations are predicted with 98.7% precision and 97.1% recall on the training set, and 95.3% precision and recall on the test set (Table 6a and Table 6b, respectively). Per abstract, an average of 96.3% of manually labelled locations are correctly returned by the GPT-4 predictions in both the validation set and the test set (Table S2.2). No-locations (i.e., no location mentioned in the abstract for the respective species) are generally predicted only marginally worse, with the exception of a notably lower precision on the training set (Table 6a), which stems from a total of 14 false negative predictions (Fig. 4). These occur across 4 abstracts, with most (9 out of 14) comprising empty predictions, and the remaining (5 out of 14) comprising predictions of insufficient information which stem from a single abstract (predictions of “North Island” rather than “Australia”).

**Table 6.**
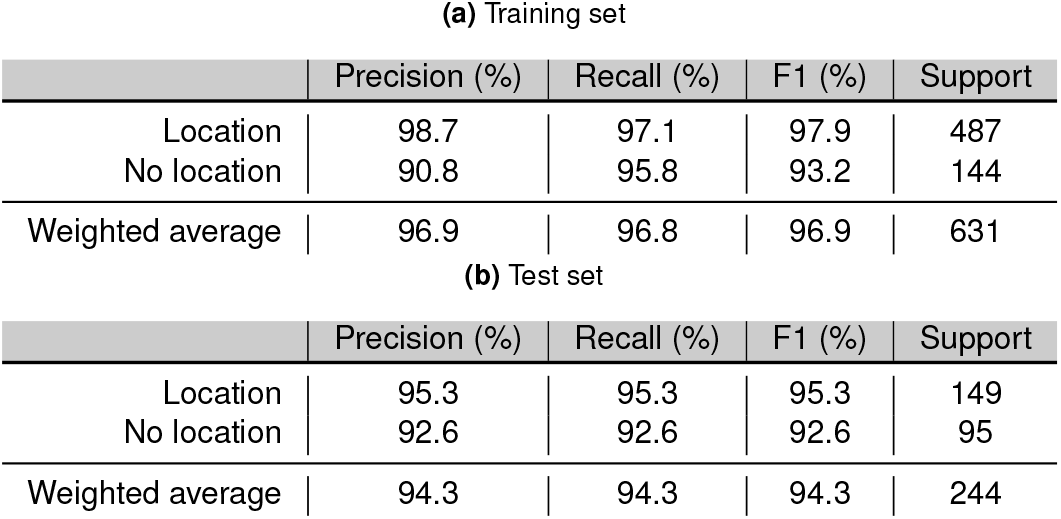
Precision and recall obtained by GPT-4 for geographic locations in the training set (a) and the held-out test set (b). Support: The number of locations and non-locations present in the manual labels. Weighted average: The average of the respective column, weighted by the support of each row. ‘Location’ refers to the location associated with the study of the species and ‘No location’ refers to the case where no location is mentioned in the abstract; If GPT-4 predicted a location that did not correspond to the manually labelled location, this is designated a false-positive, and if GPT-4 predicted no location for a manually labelled location, this is labelled as a false-negative.

| (a) Training set |  |  |  |  |
| --- | --- | --- | --- | --- |
|  | Precision (%) | Recall (%) | F1 (%) | Support |
| Location | 98.7 | 97.1 | 97.9 | 487 |
| No location | 90.8 | 95.8 | 93.2 | 144 |
| Weighted average | 96.9 | 96.8 | 96.9 | 631 |
| (b) Test set |  |  |  |  |
|  | Precision (%) | Recall (%) | F1 (%) | Support |
| Location | 95.3 | 95.3 | 95.3 | 149 |
| No location | 92.6 | 92.6 | 92.6 | 95 |
| Weighted average | 94.3 | 94.3 | 94.3 | 244 |

**Figure 4.**
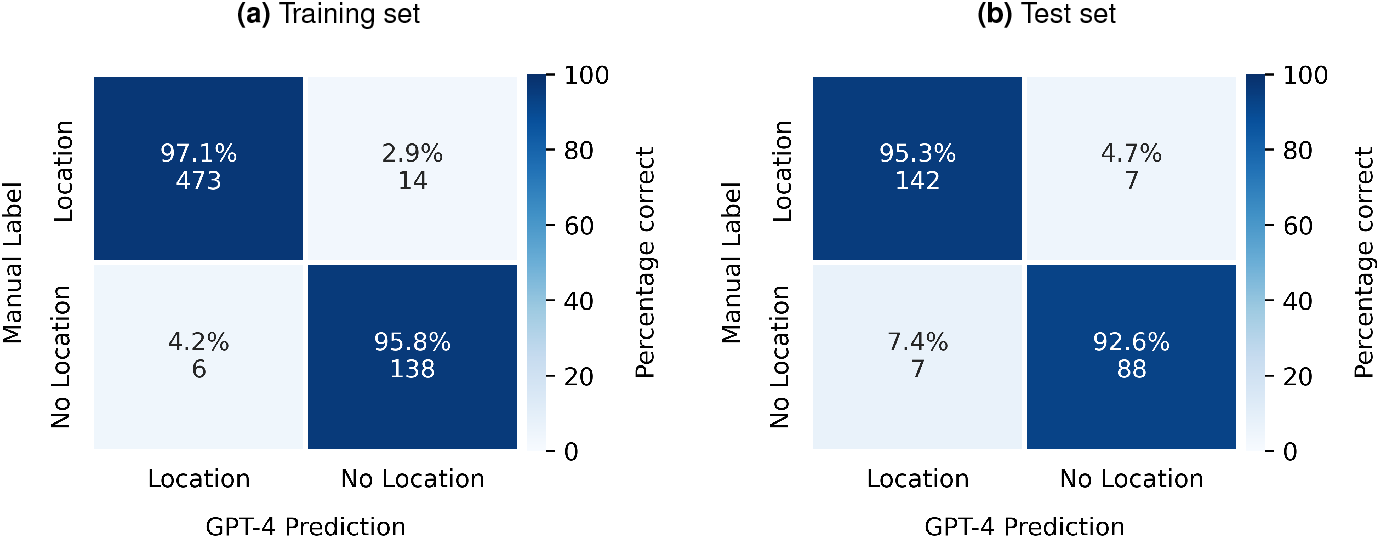
Confusion matrices of the geographic locations of the studies mentioned in the text, either as ‘Manual label’ or as ‘GPT-4 prediction’, for the training set (a) and the held-out test set (b). Values indicate the percentage of total predictions across each row (recall) and below these the corresponding absolute numbers of predictions. A true-positive prediction constitutes the prediction of a location that corresponds to, or is in close agreement with the manual label; a false-positive constitutes the prediction of a non-empty location which does not correspond to the manual label; a true-negative constitutes a location correctly left blank (i.e., no location was mentioned in the abstract); and a false-negative constitutes a location incorrectly left blank or conveying insufficient information (i.e., a location that was mentioned in the abstract but not captured by GPT-4).

## Discussion

The results presented here show that GPT-4 possesses an effective ability to (1) extract taxonomic and geographic entities from abstracts, (2) identify the roles of species from their descriptions in the abstract and (3) extract relations between entities in the abstract, such as between pest controllers and pests, and pests and certain industries as well as their affected products. However, a number of observations and caveats merit discussion.

### Hallucinations

The vast pool of information that GPT-4 is able to synthesise appears to provide a notable strength, enabling the model to recognise common usages of language in ecology such as the conventions regarding taxonomic ranks and the meaning of domain-specific vocabulary. Moreover, it allows the model to make predictions that go beyond merely the information provided in the text, allowing it to correct spelling mistakes, disentangle ambiguities, and provide appropriate roles of species based on descriptions in the text.

The ability of GPT-4 to synthesise vast quantities of data can also be a source of hallucinations, however. This refers to cases of believable, but fabricated information that have been observed to be produced by large language model chatbots such as ChatGPT (Azamfirei et al., 2023). We observed a total of 43 hallucinated species as rows in the table output of GPT-4 (all in the training set). While the first part of the prompt (Fig. 1) consistently returned the correct set of species, a subsequent mismatch was observed between this set and the species returned in the final table. This was observed exclusively in cases where GPT-4 was unable to complete the entire table in one prompt-completion and thus required multiple completions to finish the table (through the usage of a ‘continue generating’ function available in the web-interface). We suspect that this repeated generation of output may be the cause for these hallucinations, possibly as a result of a diminishment of the model’s internal memory (Gong et al., 2023) of its earlier response to the first part of the prompt. Indeed, the only cases of hallucinated species rows in GPT-4’s output (across both training and test sets) were detected in the two longest tables generated by GPT-4 (corresponding to two abstracts in the training set), consisting of 25 and 58 rows, respectively. Furthermore, all 18 species rows that were missed by GPT-4 in the training set originate from these same two abstracts, and thus the same mechanism may be responsible for these false negatives. While hallucinations are concerning, it is expected that usage of GPT-4 with higher token-lengths for prompt completion would alleviate this issue. For example, updated versions of GPT-4 are stated to have maximum token-lengths of 8,192 and 32,768 tokens respectively, while web-interface usage was limited to 2,048 tokens per generation at the time of writing (OpenAI, 2023b). The fact that the GPT-4 output for the test set contained no tables larger than 12 rows in length may thus explain why we observed no instances of hallucinated species entries in the test set.

Additionally, a small number of hallucinations were observed for the ‘Industry Type’ column in the generated data sets, where associations of pests with agriculture or forestry were stated without any basis in the text and no clear basis outside of the text. This was observed on 18 (out of 631) occasions in the training set and on 8 (out of 244) occasions in the test set. The apparent inability of GPT-4 to state whether information was drawn from inside the text or from outside the text, and if so, from where outside the text, is a well-known caveat of the model. However, new plug-ins are continuously being released to address this problem and may have utility in tasks such as demonstrated here. Connecting GPT-4 to live internet sources, as exemplified by the GPT-4-powered Microsoft Bing Chat, may also provide an effective way to reduce hallucinations through the reliable citing of sources.

The proneness of large language models to fabricate erroneous, but credible-sounding pieces of information is the subject of increasing discussions in the broader scientific community, with concerns over the accuracy, reliability and accountability of scientific output obtained with the usage of LLMs (Birhane et al., 2023). Thus, although AI models are increasingly being used in science and have begun delivering numerous scientific advances and discoveries (Wang et al., 2023), it is important for scientists to be aware of the limitations of AI tools, such as LLMs and other black-box, deep learning based models, and the potential impact of these models on reliability and reproducibility in science.

### The problem of ambiguity

The task posed to GPT-4 in this work was, furthermore, challenged by ambiguity. For example, the task of deciding whether a predator can be safely inferred to be a pest controller if it is mentioned jointly with a pest in a biological control experiment, but there is no explicit mention of predation on the pest, is highly ambiguous. So, too, is the task of deciding whether a pest is associated with agriculture in cases where there is no explicit mention of agriculture but the text includes terms often associated with agriculture such as ‘biological control’ and ‘pesticide’. Capturing such roles and associations correctly, while avoiding false positives (e.g., from nontarget species in pesticide experiments), is a difficult task that is inherently limited by both the clarity and the length of the text. Usage of LLMs such as GPT-4 for text mining tasks in ecology must be partnered with an aim to minimise these ambiguities. As LLMs become more capable of processing large prompts, definitions of terms and ‘if-then’ clauses could be more extensively included and be further refined. In the context of this work, this could consist of including a set of definitions for the various roles the model is instructed to identify and a stricter protocol on how associations are to be made (e.g., what information suffices for an “association with agriculture”?).

### Ambiguity in species roles

In the context of the species roles, the 14 labels of ‘predator’ that were not captured by GPT-4 in the training set (Fig. 3a) refer to a single abstract which mentions various species of silver fly “associated with Chamaemyiidae-based biological control programs”. Although the abstract mentions no predatory role, a predatory role was assumed during manual labelling based on the fact that silver flies are known predators – a piece of information evidently not available to GPT-4 in this instance. In other instances, however, GPT-4 does appear to have utilised information outside of the abstract to obtain its returned roles: In the case of one abstract, species that are described simply as natural enemies (and manually labelled as such) were labelled by GPT-4 as predators, which was consequently validated by other internet sources. Also in the training set, the only 4 manual labels of ‘parasitoid’ that GPT4 failed to capture include one ambiguous case (where the species in question is mentioned simply as a biological control agent but in close association with another parasitoid, and thus perhaps GPT-4’s assessment is correct), but in the 3 remaining cases a parasitoid role is clearly stated in the text, and is, in fact, reiterated by GPT-4 itself in the descriptions that it was instructed to provide (column 13; Table 1). Why it described these species as parasitoids but failed to state their roles as such is unclear. In another case, where a hyperparasitoid was labelled as a parasitoid, GPT-4 also notes the species to be hyperparasitic in its description: Why it was not labelled as such is, again, unclear.

In the test set, we observed a substantial reduction in precision and recall for predator roles as compared with the training set (Table 5). The drop in precision can be seen to be a result of 10 false positive predictions, which occur across 6 individual abstracts. One of these cases refers to a species described as an ectoparasite and manually labelled as such, but later in the abstract is described (in a German translation) as a “predatory enemy”. Not only does this example highlight the importance of consistent usage of language (e.g., the adjective ‘predatory’ to be distinct from ‘parasitic’), but it also demonstrates the capability of GPT-4 to seamlessly process text in different languages. The remaining cases refer to species described in the abstract simply as natural enemies or biological control agents but which may be inferred to be predators based on their taxonomy (e.g., spiders and beetles), or species described as entomopathogenic nematodes (EPNs), for whom it is unclear on what basis GPT-4 predicted a predatory role. The drop in recall is a result of a single abstract in which GPT-4 labelled one species as ‘prey’ and four species as ‘competitor’, rather than correctly as predators. While the error of mislabelling a predator as prey may appear substantial, the topic of this particular abstract (intra-guild predation between native and invasive spiders) provides an understandable point of confusion.

Finally, the ‘pest’ labels missed by GPT-4 in the training set comprise a total of 28 cases (across 8 abstracts), where GPT-4 chose instead to describe the species as host, prey, herbivore or left their role blank (i.e., ‘unclear’). In none of these 28 cases, the species are explicitly described as pests but were manually inferred to be acting as pests as a result of being the target of a biocontrol-related feeding trial or an insecticide experiment. The difficulty for GPT-4 appears to arise when these species are described in the text with adjectives such as ‘prey’, ‘host’ or ‘herbivore’ and the broader context in which they are mentioned in the text (e.g., as the target species of an insecticide experiment) is neglected. While we attempted to catch such cases with the prompt line “`the species must be mentioned explicitly as a pest, or at least in association with biological control, destruction, infestation, etc.`” (Fig. 1), it may be worthwhile to highlight the aforementioned cases more specifically in the prompt.

### The challenge of prompt design

The question of prompt design (or prompt engineering) comprises a serious challenge to the adoption of large language models in scientific research. Here, we proceeded to optimise the performance of the prompt design against a training set. Our optimisation approach had two main caveats. The first relates to the reliance on manual, trial-and-error based optimisation, which is time-consuming and may be intractable for very large data sets. The second relates to the problem of overfitting, which, although not well-defined in this context (as there is no proper loss function), may arise if the prompt is too tailored to the training set and fails to generalise to the rest of the data. Regardless of the optimisation strategy, however, designing the prompt to take into account the various intricacies of the task (definitions, examples, counterexamples, clarifications, reasoning) is highly non-trivial and highly data-dependent, as exemplified by the extensive prompt utilised in this work. Furthermore, performance of GPT-4 was observed to be a highly non-linear function of the prompt-design, with small changes in the prompt leading to large changes in the output (for better or for worse).

As opposed to extensive prompt engineering, a powerful alternative may be posed by the fine-tuning of the actual parameters of the model. Also known as transfer learning (Pan and Yang, 2010), fine-tuning allows users to customise the pre-trained machine learning model to their own use cases by re-training the model (or a subset of parameters of the model) on a much smaller, bespoke data set. With many applications in image and text classification (Weiss et al., 2016), transfer learning is already a rich and and active field in biomedicine, and, to a lesser degree, ecology. OpenAI has released a fine-tuning option for GPT-3.5, which is stated to “match, or even outperform, base GPT-4-level capabilities on certain narrow tasks” (Peng et al., 2023). Furthermore, fine-tuning of GPT-3.5 is stated to allow for a shortening in prompts by as much as 90% as instructions can be fine-tuned into the model itself. Fine-tuning may thus provide a promising addition, or alternative, to prompt engineering for large language models.

### Comparison with other tools

To put some of the results obtained in this work into a broader perspective, a comparison can be made with previous studies that have utilised text mining tools for applications in ecology. Previous attempts to extract taxonomic terms from abstracts have resulted in mean recall scores per abstract of 79.5% (Millard et al., 2020) and 93.6% (Cornford et al., 2022) with the help of the R package Taxize (Chamberlain and Szöcs, 2013), while the extraction of geographic locations has been achieved with a mean recall of 82.1% per abstract (Cornford et al., 2022) with the help of the CLIFFCLAVIN geoparser model (D’Ignazio et al., 2014). Furthermore, an extensive comparison of eight taxonomic named entity recognition (NER) models over four gold standard ecology corpora (Le Guillarme and Thuiller, 2022) reports scores for approximate matches ranging between 78–96% (precision), 74–93% (recall) and 76– 91% (F1-score).

Comparison with these previous studies demonstrates the capability of GPT-4 on similar tasks to be highly competitive. If we can correctly assume the risk of hallucinated species extractions to be minimised with sufficiently large token lengths, we may assume such hallucinated entries to pose a relatively low risk for future endeavours, which are likely to incorporate longer token lengths (both as input and for completion) as large language models develop. As such, our results on the test set, which did not suffer from hallucinated species extractions, may offer a valuable demonstration of the model’s potential performance. The extraction of species from abstracts in the test set was achieved with a total precision of 100%, recall of 99.6% and F1score of 99.8% (Table 2a). Investigating only the taxonomy of the extracted species, we report PC scores on the test set of 98.3–100.0% for taxonomy that was stated in the abstract (Table 3) and 96.7–99.4% for taxonomy that was not stated in the abstract (Table 4). Geographic locations (as approximate matches) were extracted from the test set with a precision, recall and F1-score of 95.3% (Table 6b), which is only marginally worse than the performance obtained from the training set. We highlight that GPT-4 achieved these results without prior training on these tasks, while simultaneously generating answers to multiple other tasks laid out in the prompt, such as extracting appropriate species roles, identifying pest controllers, pest names and associations, and providing thorough descriptions.

## Conclusion

In this work, we explored the potential of the next generation of large language models for the automation of knowledge synthesis in ecology through the application of GPT-4 to a body of literature on pest control. To this end, we prompted GPT-4 using a set of instructions involving the extraction of species, taxonomy and geographical locations, the labelling of roles, recognition of pest-controlling behaviour, pest types, pest associations and mutual relations.

The results show that GPT-4 is highly capable of this task, with performance on all investigated sub-tasks largely congruent with the manual labels. Some of the discrepancy appears to be a result of a certain degree of ambiguity in the abstracts and in the task itself. We thus restrained from speaking of ground-truth in this work, since even manual labels are subject to ambiguity and human error: Indeed, we observed a number of cases where the predictions of GPT-4 were more accurate than the manual labels. That being said, we also observed cases of hallucinations, i.e. fabricated information. In the case of hallucinated species entries in the generated tables, this appears to be a symptom of the limits (in terms of token-lengths) placed on the prompt completion. In the case of individual pieces of fabricated information, it is more difficult to identify root causes as they may be more dependent on model parameters and the original training data of GPT-4.

We hope that this work makes a valuable contribution to the swiftly evolving domain of the automation of knowledge syntheses in ecology by demonstrating the potential of the general-purpose LLM GPT-4 for such tasks. Indeed, general-purpose LLMs may provide an interesting way forward in ecology, since there is currently a lack of specialised, pre-trained language models for use in this domain. Combined with tailored prompt engineering, such models can be used for a broad range of tasks related to the extraction of information from text, and have the potential to save a large amount of time spent on manual labelling. Through their vast information base, these models can, in principle, be applied to literature spanning myriad languages, helping to mitigate the widespread bias of the English language in knowledge syntheses (Konno et al., 2020; Amano et al., 2021) although current LLM performance on underrepresented languages has been found to be poor (Laskar et al., 2023). Furthermore, their reliability may be boosted through the integration of live internet access and a growing number of plug-ins that address the traceability of information. Additionally, an ability to handle the full-texts of papers, rather than just abstracts, is likely to be tractable in the near future and improve the reliability of extracted information.

Finally, we reiterate that although we investigated the application of GPT-4 to an instance of knowledge synthesis of biological pest control, the conclusions of this work are to be understood in the larger context of the potential of general-purpose LLMs for the automation of knowledge syntheses in ecology. In the spirit of open science, such applications ideally transpire in an open-source context, ensuring both open access and reproducibility. While GPT-4 does not currently adhere to such standards, we have nonetheless aimed in this study to provide insight into the current state-of-the-art of LLMs and their potential use for automated knowledge synthesis in ecology.

## Code and data availabilit

All code and data needed to generate the figures in this paper are available on Github: https://github.com/dscheepens/GPT4-pest-controllers.

## Author contributions

Daan Scheepens and Tim Newbold conceived the ideas and designed methodology; Daan Scheepens collected the data and carried out the manual labelling, analysed the data and led the writing of the manuscript. All authors contributed critically to the drafts and gave final approval for publication.

## Bibliography

Akella, L. M., Norton, C. N., and Miller, H. (2012). Netineti: discovery of scientific names from text using machine learning methods. BMC Bioinformatics, 13(1):211. 10.1186/1471-2105-13-211.

Almond, R., Grooten, M., Juffe Bignoli, D., and Petersen, T. Living planet report 2022 – building a nature-positive society, (2022). WWF, Gland, Switzerland.

Amano, T., Berdejo-Espinola, V., Christie, A. P., et al. (2021). Tapping into non-english-language science for the conservation of global biodiversity. PLOS Biology, 19(10): e3001296.#x2013;.

Ananiadou, S., Rea, B., Okazaki, N., et al. (2009). Supporting systematic reviews using text mining. Social Science Computer Review - SOC SCI COMPUT REV, 27:509–523. 10.1177/0894439309332293.

Anderson, S. C., Elsen, P. R., Hughes, B. B., et al. (2021). Trends in ecology and conservation over eight decades. Frontiers in Ecology and the Environment, 19(5):274–282. doi:10.1002/fee.2320.

Azamfirei, R., Kudchadkar, S. R., and Fackler, J. (2023). Large language models and the perils of their hallucinations. Critical Care, 27(1):120. 10.1186/s13054-023-04393-x.

Balzan, M. V., Bocci, G., and Moonen, A.-C. (2014). Augmenting flower trait diversity in wildflower strips to optimise the conservation of arthropod functional groups for multiple agroecosystem services. Journal of Insect Conservation, 18(4):713–728. 10.1007/s10841-014-9680-2.

Birhane, A., Kasirzadeh, A., Leslie, D., and Wachter, S. (2023). Science in the age of large language models. Nature Reviews Physics, 5(5):277–280. 10.1038/s42254-023-00581-4.

Brittain, C., Kremen, C., and Klein, A.-M. (2013). Biodiversity buffers pollination from changes in environmental conditions. Global Change Biology, 19(2):540–547. doi:10.1111/gcb.12043.

Cardoso, P., Barton, P. S., Birkhofer, K., et al. (2020). Scientists’ warning to humanity on insect extinctions. Biological Conservation, 242:108426. doi:10.1016/j.biocon.2020.108426.

Chamberlain, S. and Szöcs, E. (2013). Taxize: Taxonomic search and retrieval in r. F1000Research, 2:191. 10.12688/f1000research.2-191.v1.

Chen, Q., Sun, H., Liu, H., et al. (2023). A comprehensive benchmark study on biomedical text generation and mining with chatgpt. bioRxiv. 10.1101/2023.04.19.537463.

Cohen, A. M., Ambert, K., and McDonagh, M. (2012). Studying the potential impact of automated document classification on scheduling a systematic review update. BMC Medical Informatics and Decision Making, 12(1):33. 10.1186/1472-6947-12-33.

Cornford, R., Deinet, S., De Palma, A., et al. (2021). Fast, scalable, and automated identification of articles for biodiversity and macroecological datasets. Global Ecology and Biogeography, 30(1):339–347. doi:10.1111/geb.13219.

Cornford, R., Millard, J., González-Suárez, M., et al. (2022). Automated synthesis of biodiversity knowledge requires better tools and standardised research output. Ecography, 2022(3):e06068. doi:10.1111/ecog.06068.

Dainese, M., Martin, E. A., Aizen, M. A., et al. (2019). A global synthesis reveals biodiversity-mediated benefits for crop production. Science Advances, 5(10):eaax0121. 10.1126/sciadv.aax0121.

Dey, N., Gosal, G. Zhiming, et al. Cerebras-gpt: Open compute-optimal language models trained on the cerebras wafer-scale cluster, (2023).

Dornelas, M., Antão, L. H., Moyes, F., et al. (2018). Biotime: A database of biodiversity time series for the anthropocene. Global Ecology and Biogeography, 27(7):760–786. doi:10.1111/geb.12729.

D’Ignazio, C., Bhargava, R., and Zuckerman, E. Cliff-clavin : Determining geographic focus for news articles [extended abstract]. (2014). URL https://api.semanticscholar.org/CorpusID:31483241.

Farrell, M. J., Brierley, L., Willoughby, A., et al. (2022). Past and future uses of text mining in ecology and evolution. Proceedings of the Royal Society B: Biological Sciences, 289 (1975):20212721. 10.1098/rspb.2021.2721.

Fink, M. A., Bischoff, A., Fink, C. A., et al. (2023). Potential of chatgpt and gpt-4 for data mining of free-text ct reports on lung cancer. Radiology, 308(3):e231362. 10.1148/radiol.231362. PMID: 37724963.

GBIF. Gbif home page, (2023). URL https://www.gbif.org.

Gerner, M., Nenadic, G., and Bergman, C. M. (2010). Linnaeus: A species name identification system for biomedical literature. BMC Bioinformatics, 11(1):85. 10.1186/1471-2105-11-85.

Gong, D., Wan, X., and Wang, D. Working memory capacity of chatgpt: An empirical study, (2023).

Grass, I., Albrecht, J., Jauker, F., et al. (2016). Much more than bees—wildflower plantings support highly diverse flower-visitor communities from complex to structurally simple agricultural landscapes. Agriculture, Ecosystems & Environment, 225:45–53. doi:10.1016/j.agee.2016.04.001.

Gunstone, T., Cornelisse, T., Klein, K., et al. (2021). Pesticides and soil invertebrates: A hazard assessment. Frontiers in Environmental Science, 9. 10.3389/fenvs.2021.643847.

Gurr, G., Lu, Z., Zheng, X., et al. (2016). Multi-country evidence that crop diversification promotes ecological intensification of agriculture. Nature Plants, 2:16014. 10.1038/nplants.2016.14.

Hu, Y., Ameer, I., Zuo, X., et al. Zero-shot clinical entity recognition using chatgpt, (2023).

Hudson, L. N., Newbold, T., Contu, S., et al. (2017). The database of the predicts (projecting responses of ecological diversity in changing terrestrial systems) project. Ecology and Evolution, 7(1):145–188. doi:10.1002/ece3.2579.

IPBES. Global assessment report on biodiversity and ecosystem services of the Intergovernmental Science-Policy Platform on Biodiversity and Ecosystem Services, (2019). URL 10.5281/zenodo.6417333.

IPCC. Sections, pages 35–115. IPCC, Geneva, Switzerland, (2023). 10.59327/IPCC/AR6-9789291691647.

Jumper, J., Evans, R., Pritzel, A., et al. (2021). Highly accurate protein structure prediction with alphafold. Nature, 596(7873):583–589. 10.1038/s41586-021-03819-2.

Kojima, T., Gu, S. S., Reid, M., et al. Large language models are zero-shot reasoners, (2023).

Konno, K., Akasaka, M., Koshida, C., et al. (2020). Ignoring non-english-language studies may bias ecological meta-analyses. Ecology and Evolution, 10(13):6373–6384. doi:10.1002/ece3.6368.

Laskar, M. T. R., Bari, M. S., Rahman, M., et al. A systematic study and comprehensive evaluation of chatgpt on benchmark datasets, (2023).

Le Guillarme, N. and Thuiller, W. (2022). Taxonerd: Deep neural models for the recognition of taxonomic entities in the ecological and evolutionary literature. Methods in Ecology and Evolution, 13(3):625–641. doi:10.1111/2041-210X.13778.

Letourneau, D. K. Integrated Pest Management – Outbreaks Prevented, Delayed, or Facilitated?, chapter 18, pages 371–394. John Wiley & Sons, Ltd, (2012). ISBN 9781118295205. doi:10.1002/9781118295205.ch18. URL https://onlinelibrary.wiley.com/doi/abs/10.1002/9781118295205.ch18.

Letourneau, D. K., Armbrecht, I., Rivera, B. S., et al. (2011). Does plant diversity benefit agroecosystems? a synthetic review. Ecological Applications, 21(1):9–21. doi:10.1890/09-2026.1.

Li, Y., Lin, Z., Zhang, S., et al. Making large language models better reasoners with step-aware verifier, (2023).

aLPI. Living planet index, (2024). URL www.livingplanetindex.org/.

Martin, E. A., Feit, B., Requier, F., et al. Chapter three - assessing the resilience of biodiversity-driven functions in agroecosystems under environmental change. In Bohan, D. A. and Dumbrell, A. J., editors, Resilience in Complex Socio-ecological Systems, volume 60 of Advances in Ecological Research, pages 59–123. Academic Press, (2019). doi:10.1016/bs.aecr.2019.02.003. URL https://www.sciencedirect.com/science/article/pii/S0065250419300030.

McCallen, E., Knott, J., Nunez-Mir, G., et al. (2019). Trends in ecology: shifts in ecological research themes over the past four decades. Frontiers in Ecology and the Environment, 17(2):109–116. doi:10.1002/fee.1993.

Millard, J. W., Freeman, R., and Newbold, T. (2020). Text-analysis reveals taxonomic and geographic disparities in animal pollination literature. Ecography, 43(1):44–59. doi:10.1111/ecog.04532.

Oliver, T. H., Isaac, N. J. B., August, T. A., et al. (2015). Declining resilience of ecosystem functions under biodiversity loss. Nature Communications, 6(1):10122. 10.1038/ncomms10122. OpenAI. Gpt-4 technical report, (2023).

OpenAI. Models documentation, (2023). URL https://platform.openai.com/docs/models/gpt-4.

OpenAI. Tokenizer, (2023). URL https://platform.openai.com/tokenizer.

Ouyang, L., Wu, J., Jiang, X., et al. Training language models to follow instructions with human feedback. In Koyejo, S., Mohamed, S., Agarwal, A., et al., editors, Advances in Neural Information Processing Systems, volume 35, pages 27730–27744. Curran Associates, Inc., (2022). URL https://proceedings.neurips.cc/paper_files/paper/2022/file/b1efde53be364a73914f58805a001731-Paper-Conference.pdf.

Pan, S. J. and Yang, Q. (2010). A survey on transfer learning. IEEE Transactions on Knowledge and Data Engineering, 22(10):1345–1359. 10.1109/TKDE.2009.191.

Parr, C. S., Wilson, N., Leary, P., et al. (2014). The encyclopedia of life v2: Providing global access to knowledge about life on earth. Biodiversity Data Journal, 2:e1079. 10.3897/BDJ.2.e1079.

Peng, A., Wu, M., Allard, J., et al. Gpt-3.5 turbo fine-tuning and api updates, (2023). URL https://openai.com/blog/gpt-3-5-turbo-fine-tuning-and-api-updates.

Pimentel, D. (1961). Species Diversity and Insect Population Outbreaks. Annals of the Entomological Society of America, 54(1):76–86. 10.1093/aesa/54.1.76.

Sahayaraj, K. (2008). Approaching and rostrum protrusion behaviours of rhynocoris marginatus on three prey chemical cues. Bull. Insectol, 61:233–237.

Sánchez-Bayo, F. and Wyckhuys, K. A. (2019). Worldwide decline of the entomofauna: A review of its drivers. Biological Conservation, 232:8–27. doi:10.1016/j.biocon.2019.01.020.

Tahvanainen, J. O. and Root, R. B. (1972). The influence of vegetational diversity on the population ecology of a specialized herbivore, phyllotreta cruciferae (coleoptera: Chrysomelidae). Oecologia, 10(4):321–346. 10.1007/BF00345736.

Touvron, H., Martin, L., Stone, K., et al. Llama 2: Open foundation and fine-tuned chat models, (2023).

Vaswani, A., Shazeer, N., Parmar, N., et al. Attention is all you need, (2023).

Wagner, D. L., Grames, E. M., Forister, M. L., et al. (2021). Insect decline in the anthropocene: Death by a thousand cuts. Proceedings of the National Academy of Sciences, 118(2):e2023989118. 10.1073/pnas.2023989118.

Wang, X., Wei, J., Schuurmans, D., et al. Self-consistency improves chain of thought reasoning in language models, (2023).

Wei, J., Wang, X., Schuurmans, D., et al. Chain-of-thought prompting elicits reasoning in large language models, (2023).

Weiss, K., Khoshgoftaar, T. M., and Wang, D. (2016). A survey of transfer learning. Journal of Big Data, 3(1):9. 10.1186/s40537-016-0043-6.

Wratten, S. D., Gillespie, M., Decourtye, A., et al. (2012). Pollinator habitat enhancement: Benefits to other ecosystem services. Agriculture, Ecosystems & Environment, 159:112–122. doi:10.1016/j.agee.2012.06.020.

Zhang, S., Roller, S., Goyal, N., et al. Opt: Open pre-trained transformer language models, (2022).

Zhao, B., Jin, W., Ser, J. D., and Yang, G. Chatagri: Exploring potentials of chatgpt on cross-linguistic agricultural text classification, (2023).

Zhou, D., Schärli, N., Hou, L., et al. Least-to-most prompting enables complex reasoning in large language models, (2023).

