## Supplementary Information for "Large language models help facilitate the automated synthesis of information on potential pest controllers"

#### Supplementary Note 1: Figures

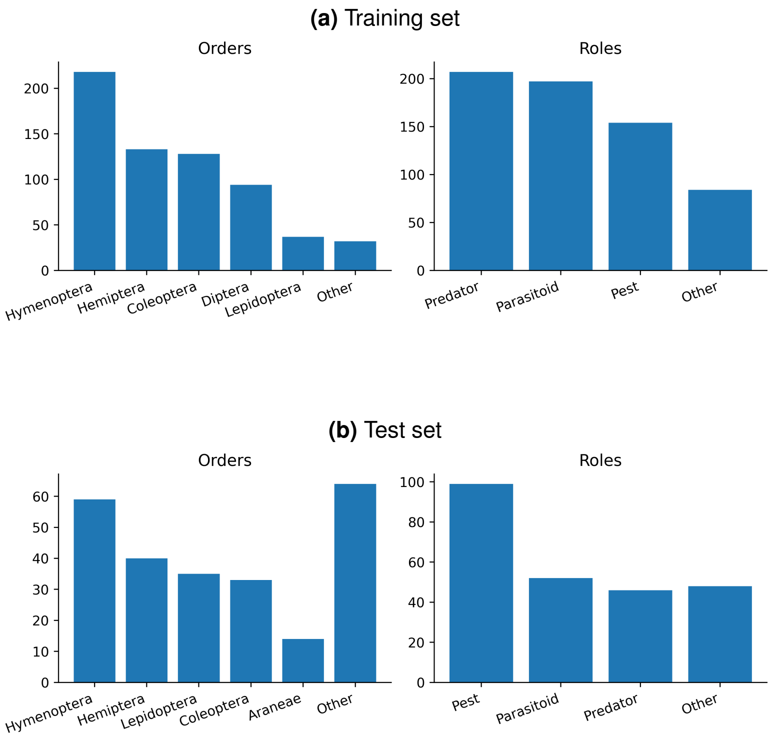

**Figure S1.1.** Occurrences (mention of a single term in a single abstract) of taxonomic orders and ascribed roles in the (manually labelled) training set (a) and the held-out test set (b). Both data sets comprise 100 abstracts.

**Title: Halotolerance of the oyster predator, Imogine mcgrathi, a stylochid flatworm from Port Stephens, New South Wales, Australia**

Abstract: The stylochid flatworm, Imogine mcgrathi was confirmed as a predator of the pteriid oyster Pinctada imbricata. Occurring at an average of 3.2 per oyster spat collector bag, the flatworms were found to consume oysters at a rate of 0.035-0.057 d-1 in laboratory trials. Predation was affected by flatworm size with larger worms capable of consuming larger oysters and of consuming greater dry weights of oyster flesh. Irrespective of flatworm size, predation was generally confined to oysters less than 40 mm in shell height. Although all predation occurred at night, shading flatworms during the day did not significantly increase the rate of predation, but there were significant increases in the dry weight of oyster meat consumed. As a means of controlling flatworm infestations, salt, brine baths (250 g kg-1) and freshwater baths were effective in killing I. mcgrathi. The ease of use of hyperor hyposaline baths then encouraged assessments of I. mcgrathi halotolerance. The flatworms were exposed to solutions ranging in salinity from 0 to 250 g kg-1 for periods of from 5 min to 3 h. Despite showing both behavioural and physiological signs of stress, I. mcgrathi survived the maximum exposure time of 3 h at salinities in the range 7.5-60 g kg-1, inclusive. Beyond this range, the duration of exposure tolerated by flatworms decreased until 0 and 250 g kg-1, at which the flatworms no longer survived the minimum tested exposure of 5 min. Thus, despite the significant impact of other stylochids on commercial bivalves, at their current prevalence, I. mcgrathi can be controlled by exposing them to hyper- and hyposaline baths for the culture of P. imbricata in Port Stephens, NSW, Australia.

Keywords: Flatworm; Halotolerance; Imogine mcgrathi; Oyster; Oyster leeches; Pinctada imbricata; Predation; Salinity; Stylochid

| Class | Order | Family | Genus | Species | Role | Generalist /Specialist | Pest Controller | Pest Names | Pest Type | Associated With | Affects | Description | Location |
|---|---|---|---|---|---|---|---|---|---|---|---|---|---|
| Turbellaria | Polycladida | Stylochidae | Imogine | mcgrathi | Predator | | TRUE | P. imbricata | Invertebrate | Aquaculture | Oysters | A stylochid flatworm that preys on the pteriid oyster P. imbricata. Can be controlled by hyper- and hyposaline baths. | Port Stephens, NSW, Australia |

**Figure S1.2.** Table generated by GPT-4 for an exemplar abstract (title, abstract and keywords) in the test set, using the fine-tuned prompt design. This is an abstract discussing a predatory flatworm affecting oyster cultures (**?**). Although the text describes Imogine mcgrathi as a predator, a broader understanding of the text reveals that the flatworm is more appropriately to be understood as acting as a pest, "infesting" oyster aqua-cultures and discussed as needing to be "controlled". In this case, GPT-4 is unable to distinguish between this nuance and incorrectly labels I. mcgrathi as a predator which controls the 'pest' oyster P. imbricata. Due to our manual limitation of pests to be either invertebrate or plant pests, this oyster is then confusingly categorised as an invertebrate pest. The oyster is also erroneously stated to affect "oysters". Clearly, the ambiguous usage of the term 'predator' in this context and its overlap with a pest role proved difficult for the model to disambiguate. The geographic location, however, is clearly stated in the text and correctly extracted.

### Supplementary Note 2: Tables

**Table S2.1.** Precision, recall and f1-score obtained by GPT-4 for each role in the training set (a) and the held-out test set (b), computed as the mean (with a spread of one standard deviation) over all abstracts in which the role is either manually labelled or predicted by GPT-4. In the majority of abstracts, GPT-4 predictions are either in full agreement with a particular role, or in none, which means that precision and recall per abstract are mostly distributed at either 0 or 1. For this reason, we observe large standard deviations for many of the roles. Cases of zero-division during computation were treated as NAN values and not included in the computation. Support: The number of abstracts where the respective role is present as a manual label.

**(a)** Training set

| | Precision (%) | Recall (%) | F1-Score (%) | Support |
|---|---|---|---|---|
| **Predator** | **100.0** $\pm$ **0.0** | **98.4** $\pm$ **11.8** | 99.2 $\pm$ 6.0 | 54 |
| **Parasitoid** | **97.2** $\pm$ **16.4** | **97.1** $\pm$ **11.6** | 97.1 $\pm$ 10.0 | 36 |
| **Pest** | **96.8** $\pm$ **17.5** | **88.8** $\pm$ **30.9** | 92.6 $\pm$ 18.6 | 70 |
| B.C.A. | 71.4 $\pm$ 45.2 | 100.0 $\pm$ 0.0 | 83.3 $\pm$ 30.8 | 7 |
| H. Parasitoid | 100.0 $\pm$ 0.0 | 83.3 $\pm$ 37.3 | 90.9 $\pm$ 22.2 | 6 |
| Nat. Enemy | NA | NA | NA | 0 |
| Prey/Host | 7.4 $\pm$ 21.0 | 33.3 $\pm$ 47.1 | 12.1 $\pm$ 28.3 | 11 |
| Other | 62.5 $\pm$ 48.4 | 71.4 $\pm$ 45.2 | 66.7 $\pm$ 33.8 | 10 |
| Unclear | 37.5 $\pm$ 48.4 | 33.3 $\pm$ 47.1 | 35.3 $\pm$ 34.0 | 14 |

**(b)** Test set

| | Precision (%) | Recall (%) | F1-Score (%) | Support |
|---|---|---|---|---|
| **Predator** | **76.0** $\pm$ **42.7** | **95.6** $\pm$ **18.6** | 84.7 $\pm$ 27.5 | 19 |
| **Parasitoid** | **100.0** $\pm$ **0.0** | **98.9** $\pm$ **5.1** | 98.9 $\pm$ 5.1 | 23 |
| **Pest** | **89.9** $\pm$ **30.2** | **96.6** $\pm$ **17.2** | 93.1 $\pm$ 18.1 | 73 |
| B.C.A. | 83.3 $\pm$ 37.3 | 71.4 $\pm$ 45.2 | 76.9 $\pm$ 30.7 | 7 |
| H. Parasitoid | 100.0 $\pm$ 0.0 | 100.0 $\pm$ 0.0 | 100.0 $\pm$ 0.0 | 1 |
| Nat. Enemy | NA | 0.0 $\pm$ 0.0 | NA | 1 |
| Prey/Host | 60.0 $\pm$ 49.0 | 50.0 $\pm$ 50.0 | 54.5 $\pm$ 36.0 | 6 |
| Other | 42.9 $\pm$ 49.5 | 60.0 $\pm$ 49.0 | 50.0 $\pm$ 37.7 | 5 |
| Unclear | 50.0 $\pm$ 50.0 | 20.0 $\pm$ 40.0 | 28.6 $\pm$ 41.6 | 10 |

**Table S2.2.** Precision and recall obtained by GPT-4 for geographic locations in the training set (a) and the held-out test set (b), computed as the mean (with a spread of one standard deviation) over all abstracts in which the label (grouped as 'Location' and 'No location') is either manually labelled or predicted by GPT-4. In the majority of abstracts, GPT-4 predictions are either in full agreement with a particular location, or in none, which means that precision and recall per abstract are mostly distributed at either 0 or 1. For this reason, we observe standard deviations that are substantial in magnitude. Cases of zero-division during computation were treated as NAN values and not included in the computation. Support: The number of abstracts where the location or non-location is present as a manual label. 'Location' refers to the location associated with the study of the species, while 'No location' refers to the case where no location is mentioned in the abstract. If GPT-4 predicted a location that did not correspond to the manually labelled location, this is designated a false-positive, and if GPT-4 predicted no location for a manually labelled location, this is labelled as a false-negative.

**(a)** Training set

| | Precision (%) | Recall (%) | F1-Score (%) | Support |
|---|---|---|---|---|
| Location | 97.1 $\pm$ 16.9 | 96.3 $\pm$ 16.6 | 96.7 $\pm$ 11.8 | 67 |
| No location | 88.9 $\pm$ 31.4 | 94.1 $\pm$ 23.5 | 91.4 $\pm$ 20.0 | 34 |

**(b)** Test set

| | Precision (%) | Recall (%) | F1-Score (%) | Support |
|---|---|---|---|---|
| Location | 92.1 $\pm$ 26.1 | 96.3 $\pm$ 17.5 | 94.2 $\pm$ 16.0 | 54 |
| No location | 93.2 $\pm$ 25.2 | 89.1 $\pm$ 31.1 | 91.1 $\pm$ 20.2 | 46 |

**Table S2.3.** Analysis of mismatches between manual labels and GPT-4 predictions of taxonomic terms available in the abstract (a) and missing from the abstract (b). $N_{\text{mismatch}}$: Number of mismatches. $N_{\text{total}}$: Number of total taxonomic terms. $N_{\text{correct}}$: Number of mismatches where the GPT-4 prediction was correct. $N_{\text{minor}}$: Number of minor mistakes made by GPT-4. $N_{\text{major}}$: Number of major mistakes made by GPT-4.

**(a)** Available taxonomy

| | | Class | Order | Family | Genus | Species |
|---|---|---|---|---|---|---|
| Training set | $N_{\text{mismatch}}/N_{\text{total}}$ | 3/146 | 6/383 | 6/449 | 9/631 | 20/631 |
| | $N_{\text{correct}}/N_{\text{mismatch}}$ | 0/3 | 6/6 | 5/6 | 6/9 | 16/20 |
| | $N_{\text{minor}}/N_{\text{mismatch}}$ | 3/3 | 0/6 | 1/6 | 3/9 | 3/20 |
| | $N_{\text{major}}/N_{\text{mismatch}}$ | 0/3 | 0/6 | 0/6 | 0/9 | 1/20 |
| Test set | $N_{\text{mismatch}}/N_{\text{total}}$ | 0/73 | 10/117 | 2/123 | 2/244 | 12/244 |
| | $N_{\text{correct}}/N_{\text{mismatch}}$ | 0/0 | 8/10 | 2/2 | 2/2 | 11/12 |
| | $N_{\text{minor}}/N_{\text{mismatch}}$ | 0/0 | 0/10 | 0/2 | 0/2 | 0/12 |
| | $N_{\text{major}}/N_{\text{mismatch}}$ | 0/0 | 2/10 | 0/2 | 0/2 | 1/12 |

**(b)** Missing Taxonomy

| | | Class | Order | Family |
|---|---|---|---|---|
| Training set | $N_{\text{mismatch}}/N_{\text{total}}$ | 1/485 | 3/248 | 18/182 |
| | $N_{\text{correct}}/N_{\text{mismatch}}$ | 0/1 | 0/3 | 2/18 |
| | $N_{\text{minor}}/N_{\text{mismatch}}$ | 1/1 | 1/3 | 3/18 |
| | $N_{\text{major}}/N_{\text{mismatch}}$ | 0/1 | 2/3 | 13/18 |
| Test set | $N_{\text{mismatch}}/N_{\text{total}}$ | 12/171 | 4/127 | 7/121 |
| | $N_{\text{correct}}/N_{\text{mismatch}}$ | 0/12 | 1/4 | 3/7 |
| | $N_{\text{minor}}/N_{\text{mismatch}}$ | 11/12 | 1/4 | 0/7 |
| | $N_{\text{major}}/N_{\text{mismatch}}$ | 1/12 | 2/4 | 4/7 |

### Supplementary Note 3: Prompt Versions

**Prompt**

```
Here are a title, abstract and keywords:

{Text}

Based on this information, you are tasked with finding species in the text, carefully classifying them and putting them into a table. How many species do you count, neglecting species that are plants, bacteria, fungi or pathogens? For each of these species, provide a row with 13 columns.

Fill in column 1 the inferred class of the species. Fill in column 2 the inferred order of the species. Fill in column 3 the inferred family of the species. Fill in column 4 the genus name. Fill in column 5 the species name. If you suspect any misspellings in the genus or species names, proceed to correct them. If the genus is found with no particular species name, leave column 5 blank.

State in column 6 whether the species is mentioned as a 'predator', 'parasitoid', 'pest', 'seed eater', 'leaf miner', 'pollinator', or anything else. If there is any mention of a species being a generalist (or polyphagous), fill in column 7 as 'generalist'. If there is any mention of a species being a specialist (or monophagous), fill in column 7 as 'specialist'. If there is mention of neither, leave column 7 blank.

If the species is mentioned as a predator, parasitoid, natural enemy or biological control agent of any other species, list them in column 8. State in column 9 whether the species listed in column 8 are insects, molluscs, weeds or anything else. State in column 10 whether the species listed in column 8 are associated with agriculture, forestry, cattle or anything else.

If the species listed in column 8 are considered pests, fill in column 11 as 'true', else 'false'. If column 11 is true, list in column 12 what plants, products or other things the pest species are a pest of. If column 6 is a 'pest', then leave columns 7 and 8 blank but fill in columns 9, 10 and 12 with reference to this pest species.

In column 13, give a thorough  justification of the assessment of columns 6 to 12. Also note here any corrections you made to the genus or species name if you did so.

Provide only the table, nothing else, and in the table, provide only the species rows (neglect the first row with the column headings).
```

**Output**

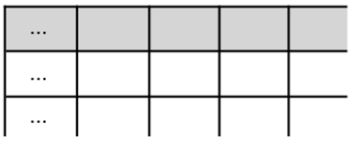

**Figure S3.1.** Prompt version 1: Initial prompt design. The prompt begins with a short outline of the task and then provides instructions on how to fill in each column in the table. We added clarifications at the end to avoid formatting issues that were observed in preliminary testing, such as entries that were shifted into the wrong column and the output of comments in addition to the table.

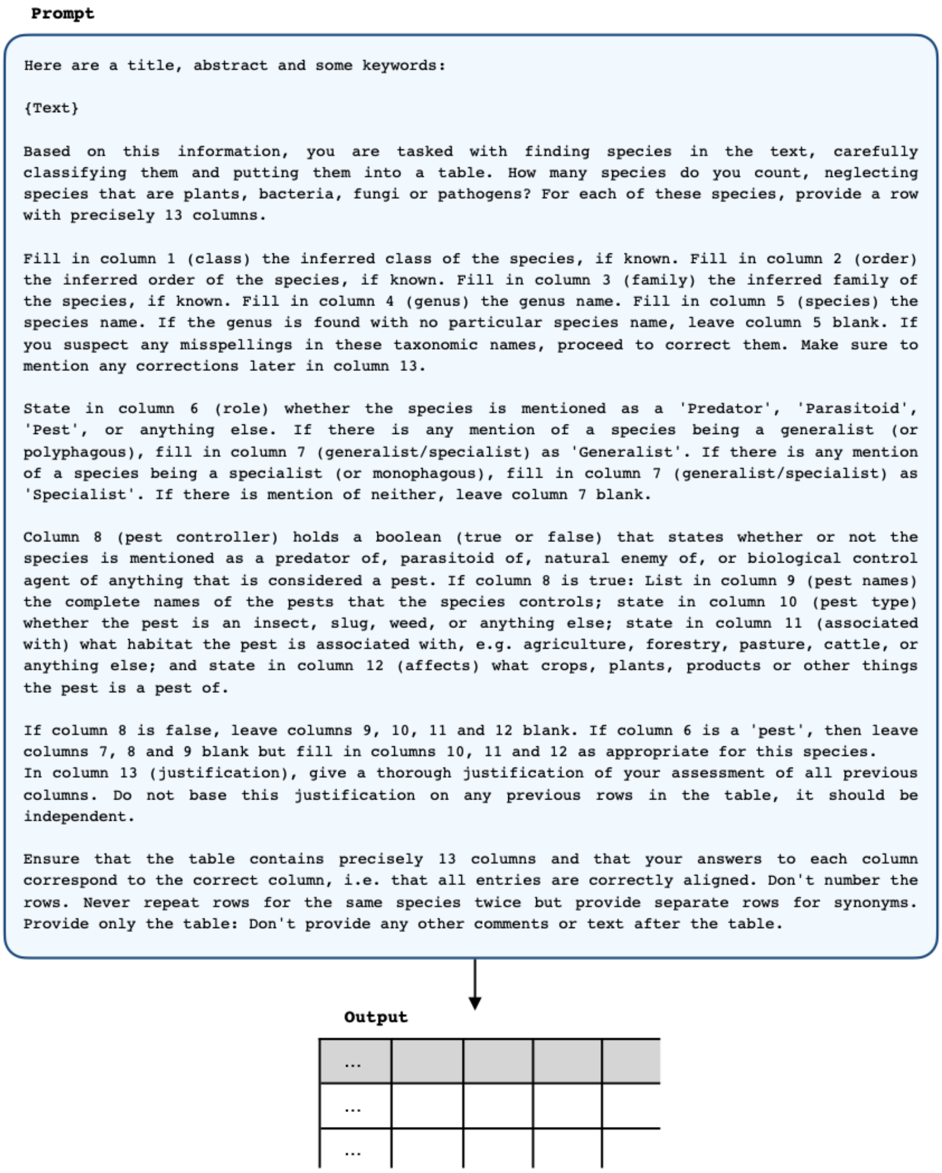

**Figure S3.2.** Prompt version 2: Prompt design following the first iteration of fine-tuning against the training set. Rather than first listing any species that are mentioned as prey or host in column 8 and then classifying these as pests or not in column 11 (as in version 1), we decided to designate a new column 'Pest Controller' as column 8 and only if this column is found to be true are the corresponding pest species to be filled in in columns 9, 10, 11 and 12. The 'Pest Controller' column allows for species to be identified as pest controllers even if no specific pests can be identified in the text. We also added the "if known" phrases for columns 1 to 3 in an attempt to reduce instances of fabricated information that we observed, and limited the examples of roles to 'predator', 'parasitoid' and 'pest' because we were getting cases where GPT-4 choose roles that were excessively specific. The additional clarifications were added at the end to attempt to avoid cases where GPT-4 numbered the rows and to ensure separate rows for each synonym.

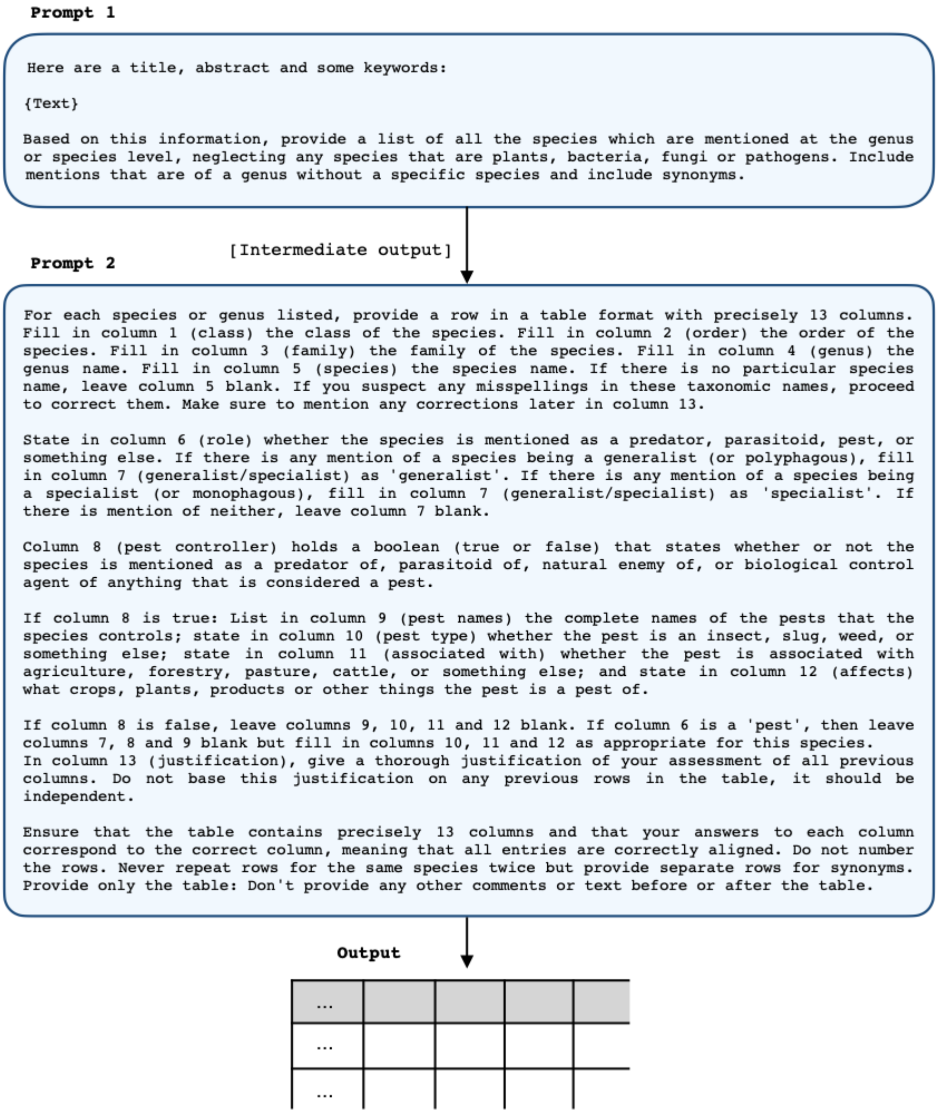

**Figure S3.3.** Prompt version 3: Prompt design attempt during the second iteration of fine-tuning against the training set. We tested out this 2-step prompt design, in which we first asked GPT-4 to list all relevant species and only then to construct a table for these species, following the idea of least-to-most prompting (**?**). The "if known" phrases from version 2 did not prove effective in avoiding mistakes in the taxonomy and were dropped.

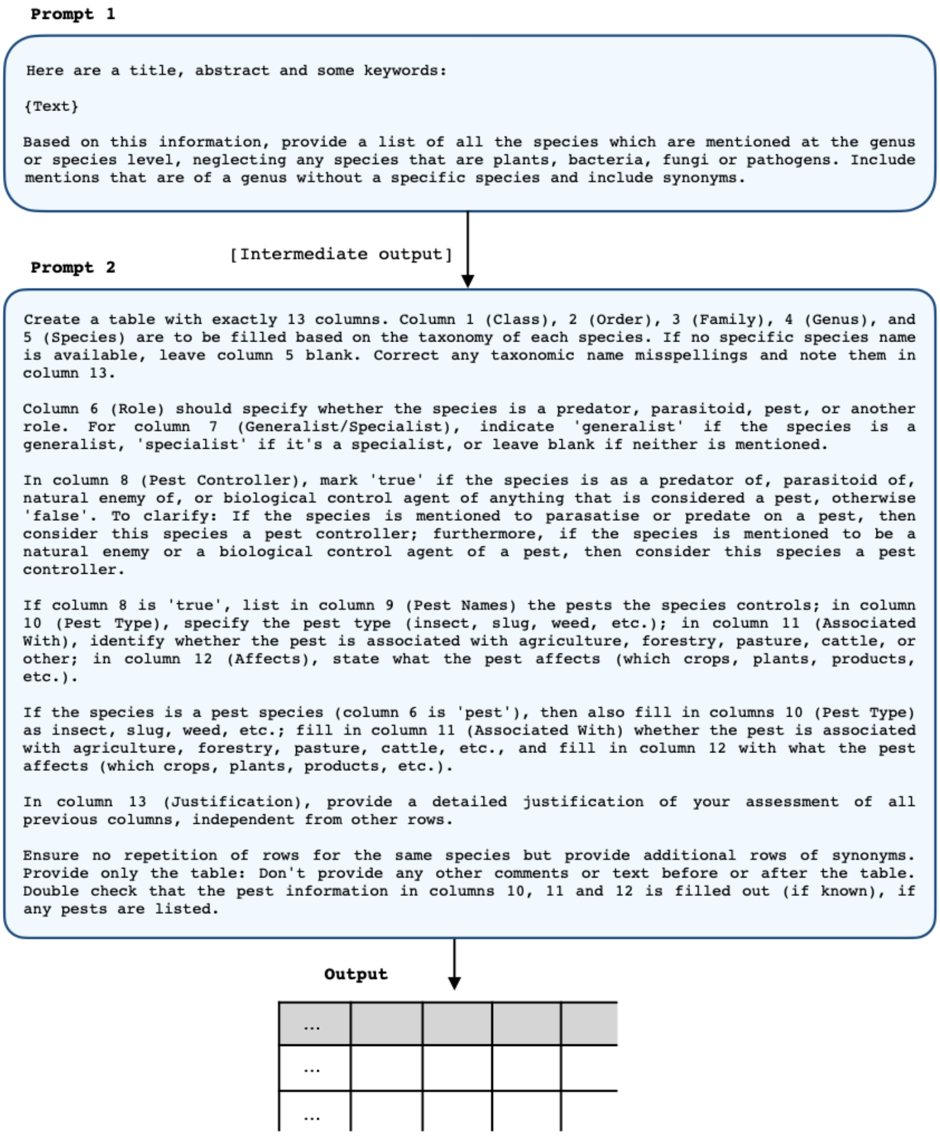

**Figure S3.4.** Prompt version 4: Prompt design attempt during the second iteration of fine-tuning against the training set. Clarifications were added for the 'Pest Controller' column and for the pest information in columns 10, 11 and 12 since these were often observed to be left out.

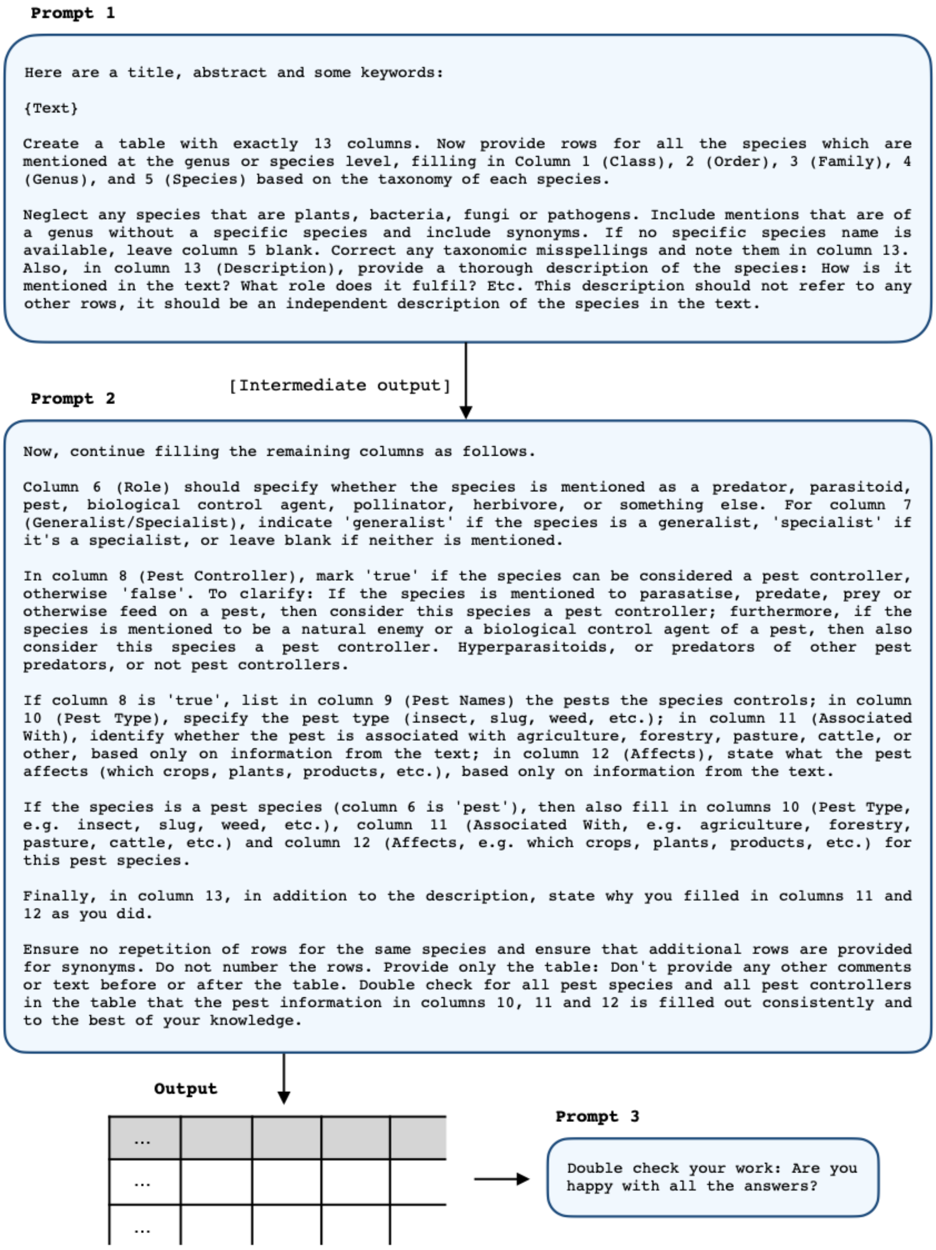

**Figure S3.5.** Prompt version 5: Prompt design following the second iteration of fine-tuning against the training set. We tested out the addition of a third prompt to investigate whether GPT-4 would be able to identify its own errors. We also tested out filling out the table in two parts and included a clarification on hyperparasitoids (parasitoids of other parasitoids) and predators of other pest predators.

**Prompt 1**

Here are a title, abstract and some keywords:

{Text}

Based on this information, provide a comprehensive list of all the species that are mentioned at the genus or species level, neglecting any species that are plants, bacteria, fungi or pathogens. Include every synonym of a species as a separate species. Double check that you have identified all species, including those that may be abbreviated (in which case, infer the full name).

[Intermediate output]

**Prompt 2**

Create a table with exactly 13 columns. Provide rows for all the species (either at the genus or species level) that you listed previously, filling in Column 1 (Class), 2 (Order), 3 (Family), 4 (Genus), and 5 (Species) based on the taxonomy of each species. If no specific species name is available, leave column 5 blank. Correct any spelling mistakes in the taxonomy. Make sure all species and genera that you listed in your previous response are included in the table.

For the remaining columns, thoroughly analyse the provided title, abstract and keywords. You are tasked with reasoning to your best ability about the way that each species is mentioned in the text. Let's think through the following tasks step by step.

For column 6 (Role), specify whether the species is mentioned as a predator, parasitoid, pest, biological control agent, pollinator, herbivore, or something else. If a species is mentioned as both a predator and a biological control agent, then choose predator, and if a species is mentioned as both a parasitoid and a biological control agent, then choose parasitoid. If a species is a prey or host of a predator a parasitoid, this does not suffice for the species to be classified as a pest; the species must be mentioned explicitly as a pest, or at least in association with biological control, destruction, infestation, etc. For column 7 (Generalist/ Specialist), indicate 'generalist' if the species is a generalist, 'specialist' if it's a specialist, or leave blank if neither is mentioned.

In column 8 (Pest Controller), mark 'true' if the species can be considered a pest controller, otherwise 'false'. To clarify: If the species is mentioned as a predator, parasitoid, natural enemy or biological control agent of (or in association with) a pest, or is mentioned to specifically prey or feed on a pest, then consider this species a pest controller; if it is stated as a biological control agent of a weed, then also consider the species a pest controller. Hyperparasitoids, or predators of other pest predators (hyperpredators), are not pest controllers.

If column 8 is 'true', list in column 9 (Pest Names) the pests the species controls; in column 10 (Pest Type), specify the pest type (insect, slug, weed); in column 11 (Associated With), identify what industry the pest is associated with (agriculture, forestry, or something else), based only on information from the text; in column 12 (Affects), state what the pest affects (which crops, plants, products, etc.), based only on information from the text.

If the species is a pest species, then fill in column 10 the pest type (insect or slug), column 11 what industry it is associated with (agriculture, forestry, etc.), and column 12 what crops, plants or other products this pest affects.

In column 13 (Description), provide a thorough description of the species: How is it mentioned in the text? What role does it fulfil? Also, justify your choices for columns 11 and 12, if filled in. This description should not refer to any other rows, it should be an independent description of the species in the text.

Do not repeat rows for the same species and ensure that additional rows are provided for synonyms. For any column: If it cannot be filled in, leave it blank. Do not number the rows: The 'Class' column must be the first column. Ensure that for all pests and pest controllers, columns 10 (Pest Type), 11 (Associated With), and 12 (Affects) are filled out consistently and accurately. If a row represents a pest, specify the pest type, the industry it is associated with, and what crops, plants or other products this pest affects. If a row represents a pest controller, list the pests it controls in column 9 (Pest Names), specify the pest type (column 10), identify what industry the pest is associated with (column 11), and state what the pest affects (column 12). If it cannot be filled in, leave it blank.

Finally, thoroughly review your work. Are all species included in the table that you listed before? Are all columns filled out correctly? Have you missed anything? Provide only the table, without additional comments or text.

**Output**

| ... |
|---|
| ... |
| ... |

**Figure S3.6.** Prompt version 6: Prompt design attempt during the third iteration of fine-tuning against the training set. We added a clarification on abbreviated species names in the first part of the prompt and added a clarification on the 'Role' column in the second part of the prompt, regarding pests and species described with more than one role. The third part of the prompt in version 5 was incorporated here in the second part of the prompt.

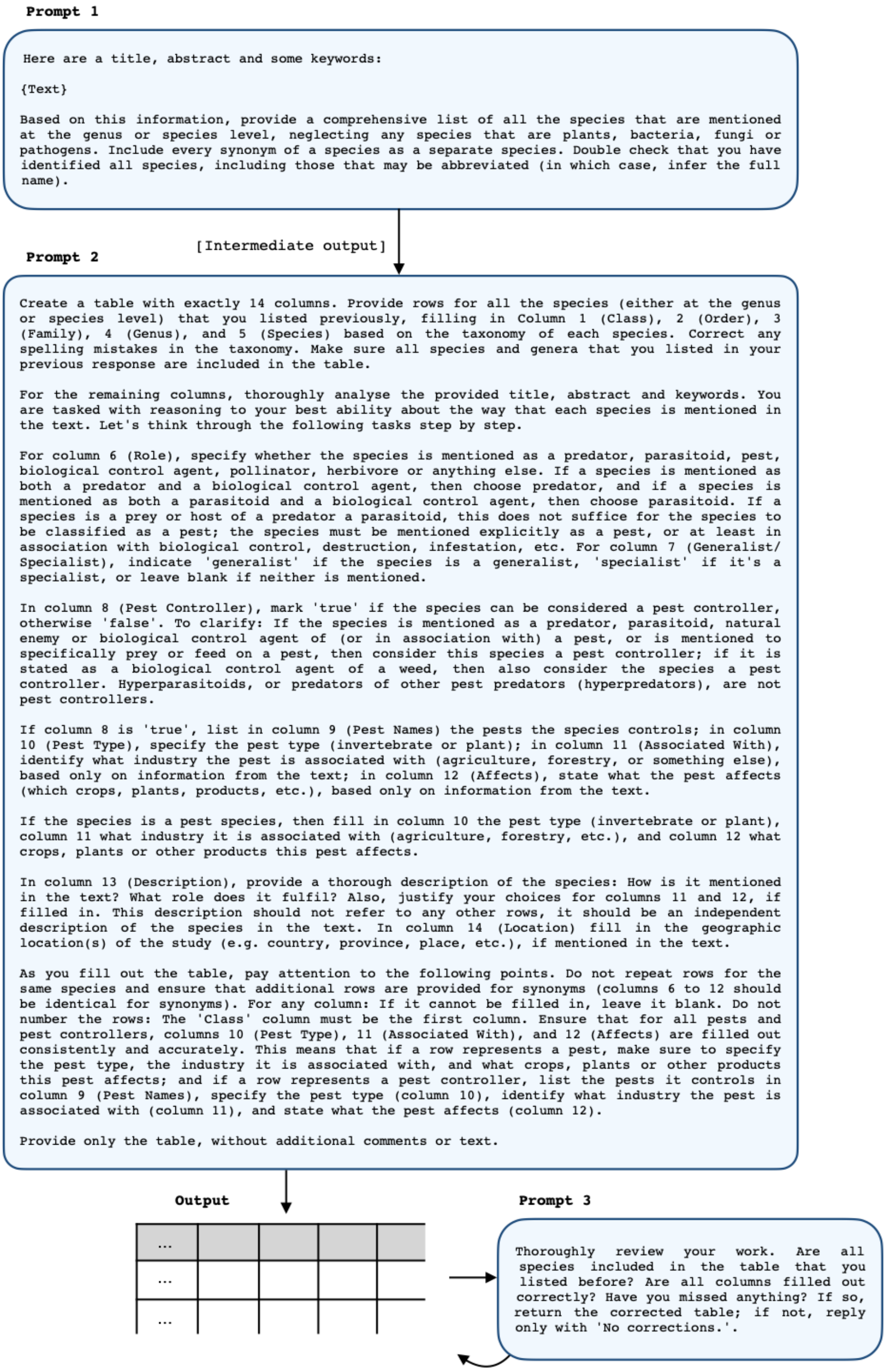

**Figure S3.7.** Prompt version 7: The final prompt design, following the third iteration of fine-tuning against the training set. We reverted back to the 3-part design, but changed the third part to prompt to return a corrected table if any mistakes were observed. We also added an extra column in which we ask for the geographical location of the study. Furthermore, the possible pest types were changed to 'invertebrate' and 'plant' in order to better distinguish pest control (of invertebrate pests) and weed control.
